# CD36 is a metabolic checkpoint for Th2 cell tissue residency during allergic airway inflammation

**DOI:** 10.1101/2025.09.17.676489

**Authors:** Anna Krone, Simon Schreiber, Nouria Jantz-Naeem, Anja Sammt, Jan Dudeck, Konstantinos Katsoulis-Dimitriou, Alexander Goihl, Oliver Hihn, Linda Black, Tobias Franz, Jonas Negele, Camilla Merten, Anna Marks, Burkhart Schraven, Martin Böttcher, Sarah Sandmann, Julian Varghese, Thomas Tüting, Dimitrios Mougiakakos, Jens Schreiber, Robert Geffers, Andreas Müller, Dirk Reinhold, Melanie Fachet, Sabine Stegemann-Koniszewski, Anne Dudeck, Bettina Weigelin, Sascha Kahlfuss

## Abstract

The prevalence of allergic diseases, including asthma, continues to rise in industrialized societies, yet the mechanisms sustaining pathogenic T helper 2 (Th2) responses remain incompletely understood. Here, we show that patients with allergic asthma exhibit elevated lipophilic volatile organic compounds in exhaled air and altered fatty acid–metabolism gene expression in sputum-derived Th2 cells. Using a mouse model of house dust mite–induced allergic airway inflammation, we find that the lipid transporter CD36 is dispensable for T follicular helper and germinal center B cell responses but is critical for maintaining lung-resident memory Th2 cells. CD36 regulates GATA3 and PPARγ expression in lung-resident memory Th2 cells and their interaction with type-2 conventional dendritic cells during airway inflammation. In human T cells, pharmacological inhibition of CD36 does not impair initial activation but blocks terminal Th2 differentiation. These findings identify CD36 as a metabolic checkpoint that sustains Th2 effector function and tissue residency, and establish lipid metabolism as a yet unrecognized therapeutic target in allergic asthma.

**Summary:** Allergic asthma is marked by rising prevalence yet the drivers of persistent T helper 2 (Th2) immunity remain unclear. We show that asthma patients exhibit altered fatty acid–metabolism signatures in sputum Th2 cells and elevated lipophilic volatile organic compounds in exhaled air. In a mouse model of house dust mite–induced airway inflammation, the lipid transporter CD36 was dispensable for germinal center responses but essential for lung-resident memory Th2 cells, controlling GATA3 and PPARγ expression and promoting cDC2 interactions. Pharmacological inhibition of CD36 in human T cells preserved activation but blocked terminal Th2 differentiation. These findings identify CD36 as a metabolic checkpoint that sustains Th2 effector function and tissue residency, and nominate it as a therapeutic target in allergic asthma.

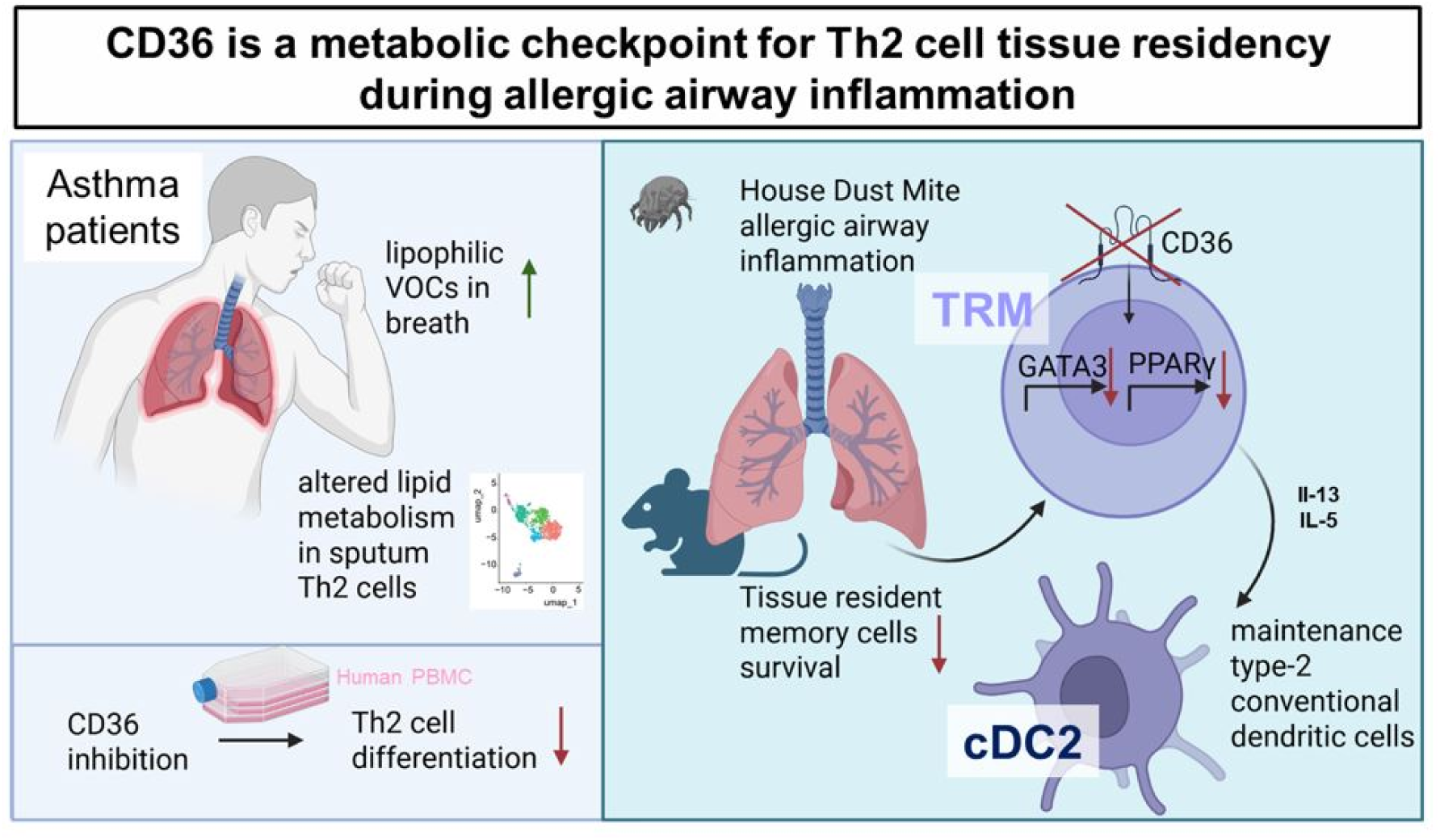

## Introduction

Asthma affects over 300 million people worldwide, and climate change is projected to further exacerbate its burden, with emergency department visits expected to rise by 8–14% by 2090, according to the Lancet 2023 report (Beggs et al., 2023; World Health Organization, 2024). The disease is characterized by airway hyperresponsiveness and inflammation, manifesting as bronchoconstriction, smooth muscle hypertrophy, goblet cell metaplasia, and ultimately structural airway remodeling (Lambrecht & Hammad, 2015; Lambrecht et al., 2019). In the mediastinal lymph nodes draining the lung, T follicular helper (Tfh) cells provide help to germinal center (GC) B cells to produce allergen-specific IgE through IL-4, IL-21 and CD40 signaling during allergic asthma. Moreover, Tfh cells have been identified as direct precursors of terminally differentiated T helper (Th)2 effector cells in the lung (Ballesteros-Tato et al., 2016; Lambrecht & Hammad, 2015). Lung Th2 cells produce cell fate-specific effector cytokines IL-4, IL-5, and IL-13, which promote IgE production (IL-4), eosinophil mobilization and activation (IL-5), and airway remodeling (IL-13). Further, terminally differentiated Th2 cells give rise to tissue-resident memory T (Trm) cells in the lungs, which persist after initial antigen exposure and thereby significantly contribute to the molecular fixation and chronification of allergic asthma (Camacho et al., 2022; Rahimi et al., 2020).

Patients suffering from allergic asthma show an increased amount of long-chain fatty acids including palmitic acid in their bronchoalveolar lavage (BAL) (Kang et al., 2014; Ravi et al., 2021; Vasseur & Guillaumond, 2022). Consistently, during house dust mite (HDM)-induced airway inflammation in mice, bronchial epithelial cells and lung Th2 cells upregulate genes involved in fatty acid metabolism (Ravi et al., 2021; Tibbitt et al., 2019). In this context, recent single cell RNA sequencing (scRNAseq) data provided further evidence that Th2 cells in the asthmatic lung depend on lipid metabolic pathways. In addition, ATAC sequencing of lung Th2 cells showed an increased accessibility of genes involved in fatty acid oxidation, including *Cd36* (Tibbitt et al., 2019). Interestingly, transcriptional analyses have revealed that tissue-resident memory T cells (Trms) use exogenous free fatty acids and oxidative metabolism to persist in tissue (Pan et al., 2017). Th2-like Trms and circulating memory Th2 cells share a core Th2 gene program but diverge in their transcriptional signatures during HDM-induced airway inflammation, most prominently by upregulating Cd36 in lung Trms (Rahimi et al., 2020). Yet, whether CD36 governs the establishment of Th2 tissue residency and thereby the chronicity of allergic airway inflammation has remained unresolved.

The scavenger receptor CD36 binds oxidized low-density lipoprotein (oxLDL) and long-chain fatty acids (Okamura et al., 2009). Beyond serving as a lipid transporter, CD36 functions as a signaling hub that couples lipid uptake to metabolic reprogramming through peroxisome proliferator–activated receptor (PPAR)γ activation. In melanoma, it was demonstrated that the lipid-rich tumor microenvironment induces CD36 expression in regulatory T cells (Tregs), supporting their metabolic adaptation, survival, and immunosuppressive function (H. Wang et al., 2020). Conversely, CD36-initiated lipid peroxidation contributes to CD8+ T cell dysfunction in tumors (S. Xu et al., 2021). However, the role of the lipid translocase and signaling receptor CD36 in the establishment of Th2 tissue residency and, thereby, the fixation of allergic airway inflammation remained enigmatic.

Here we show that patients suffering from allergic asthma exhale increased amounts of lipids and that Th2 cells in their sputum show an altered expression of genes related to fatty acid metabolism. Mice with a T cell-specific deletion of CD36 show an unaltered Tfh and GC B cell response in mediastinal lymph nodes but reduced numbers of lung Trm cells following HDM allergic airway inflammation. Mechanistically, CD36 sustains PPARγ and GATA3 expression in Trms, promoting their persistence and crosstalk with type 2 conventional dendritic cells (cDC2s), thereby perpetuating airway pathology. In human T cells, pharmacological inhibition of CD36 does not interfere with the activation but with terminal differentiation of human Th2 cells. Together, our results provide evidence that lipid signaling is a metabolic checkpoint for T cell tissue residency during Th2 allergic asthma.

## Results

### Asthma patients show altered fatty acid metabolism gene expression in sputum Th2 cells and increased amounts of lipophilic volatile organic compounds

Sputum analysis is a valuable tool in asthma management because it allows to directly assess the type and degree of airway inflammation. To dissect the immunological and metabolic signatures underlying allergic asthma, we performed high-resolution single-cell transcriptomic profiling of sputum leukocytes from three allergic asthma patients and three unrelated healthy donors (HDs) (**Table 1**). Importantly, all three asthma patients exhibited significantly elevated serum IgE levels compared to HDs, a characteristic hallmark of allergic asthma (**Figure 1A**). For single cell RNA sequencing (scRNAseq) we isolated live CD45+ cells from HDs and asthma patients by FACS sorting and subsequently performed single-cell RNA using a10x Genomics pipeline (**Figure 1B**). (**Suppl. Figure 1A, B**). After quality control, five transcriptionally distinct immune cell clusters emerged from all sputum CD45 leukocyte samples (**Figure 1C, Suppl. Figure 1A**). According to hallmark gene expression, these clusters were annotated to CD4+ T cell, CD8+ T cell, B cells, NK cells and maccrophages. Importantly, within the equal number of CD45+ leukocytes the B cell cluster appeared more prominent in asthma patients, whereas CD4+ T cells constituted a relatively higher fraction in HD due to the lower abundance of B cells. The Th2 subcluster, defined as CD3+CD4+GATA3+ cells, exhibited a profound dysregulation of asthma-related genes, with significantly reduced expression of *MX1* and *RPS26* in HDs compared to asthma patients, alongside altered expression of metabolic genes including *ACAA2* and *HACD3*, encoding the enzymes Acetyl-CoA acyltransferase 2 and Hydroxyacyl-CoA dehydratase 3, respectively (**Figure 1D** and **1E**, and **Suppl. Figure 1B**) (Loisel et al., 2016; Safran et al., 2010). Pathway analysis using the Kyoto Encyclopedia of Genes and Genomes (KEGG) revealed that differentially expressed genes (DEGs) from CD3+CD4+GATA3+ cells were enriched not only in Th1/Th2 differentiation but also in metabolic pathways such as fatty acid metabolism (**Figure 1F**). In addition to sputum scRNAseq, we sought to identify disease-specific signatures in the breath of asthma patients. To this end, we collected breath samples from HDs and asthma patients before allergen-specific provocation (bp), immediately after and 8 hours post provocation to analyze the containing volatile organic compounds (VOCs) by Proton-transfer-reaction mass spectrometry (PTR-MS) (Fachet et al., 2022) (**Figure 1G**). Here, we found a significantly increased amount of lipophilic VOCs in the exhaled breath of asthma patients compared to HDs. Among them were 2-Butanal, 1-Buten-3-yne/Butadiene, 2-methylfuran and fatty acids including acetic acid before and after provocation (**Figure 1I** and **Suppl. Figure 1C**). Together, our results indicated that sputum Th2 cells from asthma patients show an alteration in fatty acid metabolism gene expression and that asthma patients exhale increased amounts of lipophilic VOCs.

**Figure 1:**
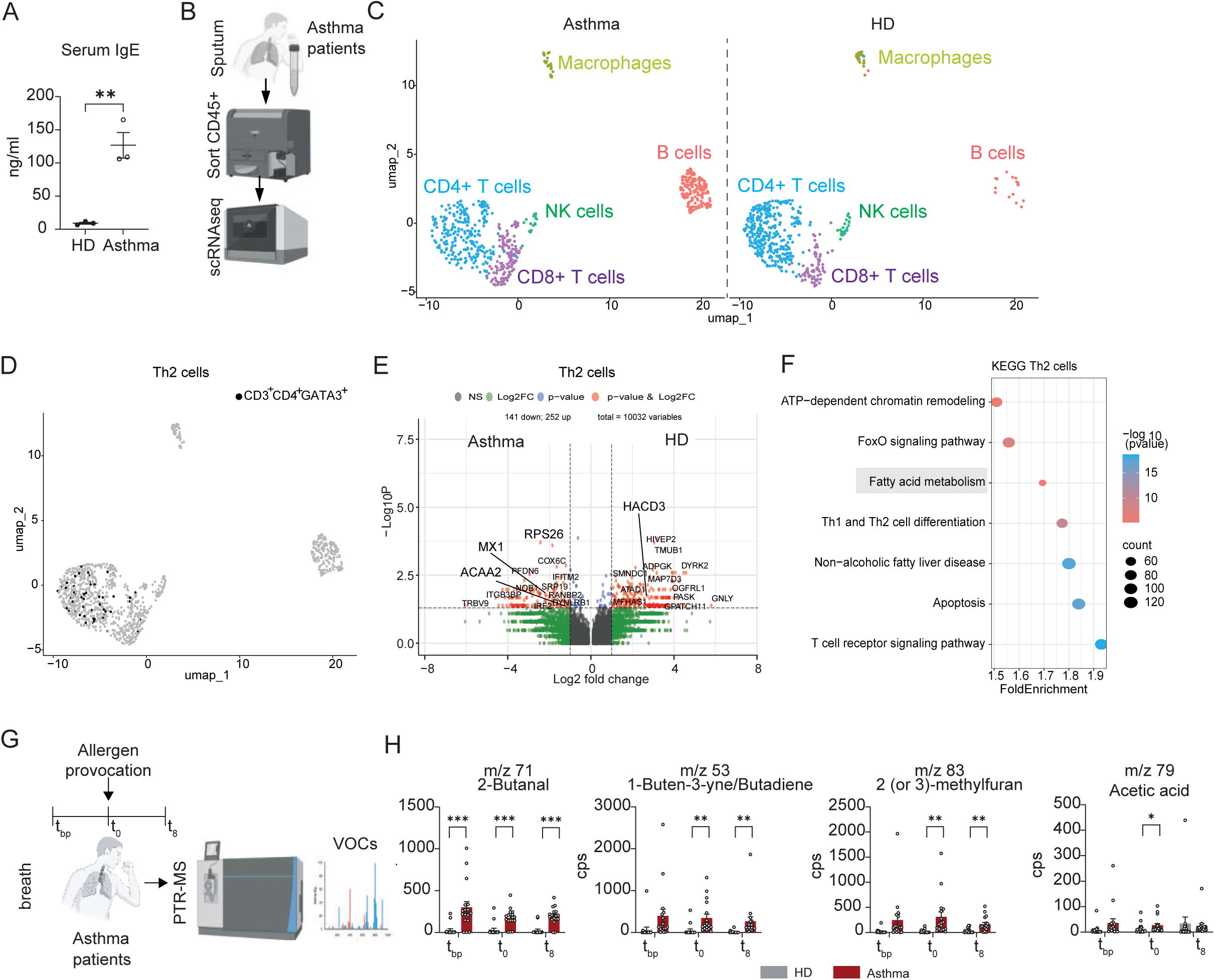
Unbiased analysis of sputum CD4+ T cells using scRNA-Seq and assessment of volatile organic compounds (VOCs) reveal significant differences in lipid metabolism in asthma. (A) Total IgE serum levels in healthy donors (HD) and asthma patients in steady state. (B) Schematic representation of sample collection and processing for scRNA-Seq from sputum samples of three healthy donors and three asthma patients. (C) Uniform Manifold Approximation and Projection (UMAP) visualization of single CD45+ sputum cells from HD and asthma patients. (D) UMAP visualization of CD3+CD4+GATA3+ subclustered Th2 cells from HD versus asthma patients. (E) Volcano plot depicting differentially expressed genes (DEGs) in CD3+CD4+GATA3^+^ subclustered Th2 cells from HD versus asthma patients. (F) KEGG pathway enrichment analysis of DEGs in the CD3+CD4+GATA3^+^ subcluster. (G) Schematic representation of breath collection from HD and asthma patients before allergen-specific provocation (bp), immediately after and 8 hours post provocation to analyze the containing volatile organic compounds (VOCs) by Proton-transfer-reaction mass spectrometry (PTR-MS) collection. (H) Lipophilic and fatty acid volatile organic compounds (VOCs) in exhaled breath from HD and asthma patients before (t_pb_), immediately after (t_0_), and 8 hours (t_8_) post sodium chlorid inhalation or allergen-specific provocation, respectively. Sample sizes: (A)-(F) HD (n = 3), asthma (n = 3); (H) HD (n = 16), asthma (n = 17). Statistical significance determined using GLM multifactorial ANOVA: *pLJ<LJ0.05, **pLJ<LJ0.01, ***pLJ<LJ0.001.

### Cd36 deletion does not affect the function or metabolic fitness of Th2 cells *in vitro*

Interestingly, recent studies have reported that asthma patients show increased amounts of lipids in their BAL (Höglund et al., 2023), although the biological meaning behind this observation remained elusive. In agreement with our own findings (see **Figure 1**), these studies demonstrated that long-chain fatty acids, such as oleic acid, rank among the most abundant lipid species in asthmatic airways. One of the most important fatty acid receptor is CD36, which can act as a lipid transporter and receptor initiating a distinct downstream signaling (Chen et al., 2022). CD36 was shown to regulate the effector function of CD8+ CTLs and Tregs in the tumor microenvironment (Ma et al., 2021; H. Wang et al., 2020; S. Xu et al., 2021). Furthermore, Tibbitt et al. demonstrated that among all lipid-related genes especially the *Cd36* gene locus was highly accessible in *ex vivo* Th2 cells isolated from the lungs of mice that underwent allergic HDM airway inflammation. In order to study the role of CD36 in Th2 cells during allergic HDM airway inflammation, we established conditional knockout mice, in which deletion of CD36 occurs under the control of the *Cd4* promoter. T cells in these mice do not express CD36 starting from the double positive (DP) stage in the thymus. Importantly, *Cd36^fl/fl^Cd4*-Cre mice showed a similar appearance, body weight, and cellular composition in thymus and spleen compared to their wildtype (WT) littermates (**Suppl. Figure 2A-C**). Specifically, we did not detect alterations in the frequencies or cell counts of double negative (DN) stages 1-4, CD4+CD8+ DP, CD4+ single positive (SP), or CD8+ SP thymocyte subsets (**Suppl. Figure 2D-G**). Further, the steady state frequencies and counts of splenic CD4+ and CD8+ T cells, B cells, and Tregs appeared inconspicuous when compared to WT control mice (**Suppl. Figure 2H-O**). To analyze whether CD36 affects the *in vitro* effector function of Th2 cells, we isolated WT and CD36-deficient CD4+ T cells, stimulated them with anti-CD3/CD28 antibodies and differentiated them into Th1 and Th2 cells in the absence or presence of oxidized Low-Density Lipoprotein (oxLDL), which was recently reported to act as a ligand for CD36 in T cells (S. Xu et al., 2021) (**Suppl. Figure 3A and 3B**). Across all conditions, CD36-deficient and WT T cells displayed comparable proliferation, apoptosis rates, and activation (**Suppl. Figure 3C-E**). In addition, activated WT and CD36-deficient Th1 and Th2 cells in the absence and presence of oxLDL showed comparable Th1 and Th2 cytokine production (**Suppl. Figure 4**), oxidative phosphorylation and glycolysis *in vitro* (**Suppl. Figure 5**). We further performed western blot analysis of the five electron transport chain (ETC) complexes in activated and differentiated WT and CD36-deficient Th2 cells in the absence and presence of Dipalmitoylphosphatidylcholine (DPPC), the main constituent of pulmonary surfactant, or in the presence and absence of the 16-carbon fatty acid palmitate. Results indicated no significant difference in ETC complex expression between WT and CD36-deficient Th2 cells (**Suppl. Figure 6A**). In addition, western blot analysis of metabolic upstream regulator proteins showed a tendency towards increased phosphorylated AMPK (pAMPK) (Thr172) and phosphorylated mTOR (pmTOR) (Ser2428) expression in CD36-deficient compared to WT Th2 cells without reaching statistical significance (**Suppl. Figure 6B**). We further detected no significant differences between WT and CD36-deficient Th2 cells for the cytosolic expression of PPARLJ, Blimp1, NFATc1 (NFAT2) and Lipin1 (**Suppl. Figure 6C**). Taken together, these data demonstrate that T cell–specific deletion of CD36 neither perturbs T cell development at steady state nor impairs in vitro Th2 cell metabolism or effector function, thereby reinforcing previous findings in CD8+ CTLs and Tregs (H. Wang et al., 2020 and S. Xu et al., 2021).

### T cell-specific deletion of CD36 does not affect T follicular helper cell-mediated IgE production by germinal center B cells during house dust mite allergic airway inflammation

Our results indicated that CD36 is dispensable for the metabolism and effector function of Th2 cells *in vitro*. However, whether CD36 is important for the function of T cell-mediated allergic airway inflammation *in vivo* so far remained elusive. To investigate the role of CD36 in T cells during-allergic airway inflammation, we employed a two-week model of HDM induced allergic airway inflammation. To this end, WT and *Cd36^fl/fl^Cd4*-Cre mice were intranasally (i.n.) sensitized with HDM extract on days 0–2 and subsequently rechallenged i.n. on days 11–14 (**Figure 2A**). At day 14, WT and *Cd36^fl/fl^Cd4*-Cre mice showed significantly elevated peribronchial airway inflammation, increased mucus production and significantly increased frequencies and cell counts of eosinophils in their BAL (**Suppl. Figure 7A-C**). LEGENDplex^TM^ analysis of the BAL did not reveal significant differences in cytokine and chemokine levels between *Cd36^fl/fl^Cd4*-Cre and WT mice (**Suppl. Figure 7D and 7E**). However, we detected an altered cytokine and chemokine composition in the serum of *Cd36^fl/fl^Cd4*-Cre compared to WT littermates (**Suppl. Figure 8A and 8B**). Specifically, we observed significantly reduced levels of BLC (CXCL13), IFNγ, TNFα, and IL-10 in the serum of *Cd36^fl/fl^Cd4*-Cre compared to WT control mice.

**Figure 2:**
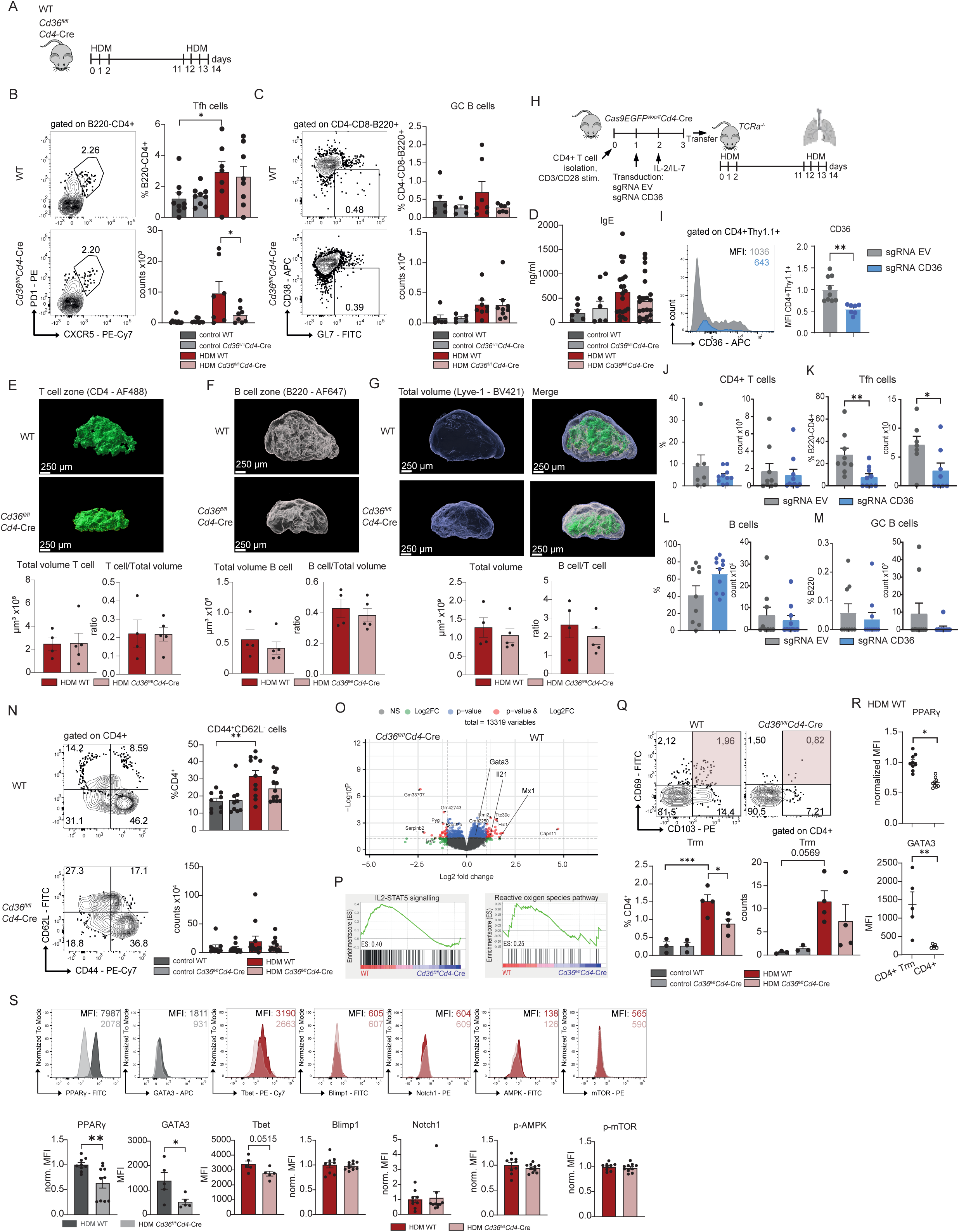
CD4 specific CD36 deletion in house dust mite (HDM) induced allergic airway inflammation does not affect cell frequencies in mediastinal lymphnodes but impairs the development of tissue resident memory (Trm) cells in the lung by reducing PPAR_γ_ and GATA3. (A) Mouse model of an acute Th2 asthmatic airway inflammation induced by house dust mite (HDM) extract using WT control mice and *Cd36^fl/f^ Cd4-*Cre. (B) Representative plots and graphs of counts and frequencies of T follicular helper (Tfh) cells in mediastinal lymph nodes from the HDM asthma model. (C) Representative plots and graphs of counts and frequencies of germinal center B cells in mediastinal lymph nodes from the HDM asthma model. (D) Total IgE serum levels of mice from the HDM asthma model. (E) – (G) 3D reconstruction of LN from WT control mice and *Cd36^fl/f^ Cd4-*Cre was performed in Imaris software. B220 was used for staining the B cell zones, CD4 for T cell zones and B220, CD4 and Lyve-1 for the total LN volume. The 3D reconstruction was performed by using the surface tool in Imaris with the help of machine learning. Cumulative graphs, showing the total volume of (E) T cell zones, (F) B cell zones and (G) total volume based on the analysis described in E. The ratios of T cell and B cell zones to total volume and in comparison to each other are also shown as a measure of relative size. (H) Viruses for CD36 was used to transduce CD4+ T cells from transgenic CD4-Cre and Cas9 expressing mice. Cells were adoptively transferred into TCRa-/-recipient mice upon induction of acute HDM allergic airway inflammation. (I) Mean Fluorescence Intensitiy (MFI) of CD36 and in CD4+Thy1.1 + cells (J) – (M) Representative plots and graphs of counts and frequencies of (J) CD4, (K) Tfh cells (L) B cells (M) GC B cells. (N) Representative plots and graphs of counts and frequencies of CD44+CD62L+ effector cells in lungs from the acute HDM asthma model. (O) Volcano plot depicting DEGs of WT versus *Cd36^fl/fl^Cd4*-Cre mice upon induction of HDM allergic airway inflammation. (P) Gene Set Enrichment analysis (GSEA) of WT versus *Cd36^fl/fl^Cd4*-Cre mice upon induction of chronic HDM allergic airway inflammation revealed IL-2–STAT5 pathway (MM3902) and the reactive oxygen species pathway (MM3895). (Q) Representative plots of tissue resident memory (Trm) cells in lungs from the acute HDM asthma model. (R) MFI of PPARγ and GATA3 in Trm cells and non Trm cells in lungs from WT mice from the acute HDM asthma model. (S) Representative histograms and MFIs of transcriptionfactors and signalin proteins in tissue resident memory (Trm) cells in lungs from the acute HDM asthma model. Sample sizes: (B)-(C) n = 6-9; (D) n = 6-21; (E)-(G) n = 5; (I)-(M) n = 7-10; (N) n = 5-10; (O)-(P) n = 3; (Q) = 4; (R)-(S) n = 5-10. Statistical analysis was performed using ordinary one-way ANOVA followed by Tukey’s test for multiple comparisons ((B)-(D), (N), (Q)) or Students T-Test ((E)-(M), (R)-(S)); *p < 0.05, **p < 0.01, ***p < 0.001.

Building on our findings of altered cytokine profiles in the blood, we next examined how these changes affect the primary immune response in the mediastinal lymph nodes, the lung-draining lymph nodes, during HDM-induced airway inflammation. Flow cytometry analyses revealed largely unaltered T and B cell populations as well as CD4+ T cell cytokine production in mediastinal lymph nodes of *Cd36^fl/fl^Cd4*-Cre compared to WT littermates (**Suppl. Figure 8C** and **Suppl. Figure 9**). Importantly, despite a significant reduction of total numbers of Tfh cells, frequencies and cell count of germinal center (GC) B cells remained unaltered in mediastinal lymph nodes from *Cd36^fl/fl^Cd4*-Cre compared to WT littermates (**Figure 2B and 2C**). Consistent with this, total IgE levels in the serum were not significant different between *Cd36^fl/fl^Cd4*-Cre and WT mice following HDM allergic airway inflammation (**Figure 2D**).

These results were supported by analysis of potential structural changes in mediastinal lymph nodes, utilizing whole-mount staining of entire mediastinal lymph nodes followed by tissue-clearing and 3D imaging. This analysis revealed preserved T and B cell zone architecture in *Cd36^fl/fl^Cd4*-Cre mice, indistinguishable from WT littermates (**Figure 2E-G**). To validate the findings of the genetically engineered mice with a CD4-specific deletion of CD36 we also deleted the lipid transporter and scavenger receptor CD36 by means of sgRNA-mediated CRSIPR/Cas9 gene targeting in CD4+ T cells. To this end, we isolated CD4+ T cells from CD4Cre X Cas9 C57BL/6N-Gt(ROSA)^26Sorem1(CAG-cas9*,-EGFP)Rsky^/J (*Cas9EGFP^stop/fl^Cd4Cre*) mice. In these mice, all T cells from the DP stage in the thymus express *Cas9*. Thereafter Cas9-expressing CD4 T cells were *in vitro* activated by CD3 and CD28 antibodies and transduced with sgRNAs (with Thy1.1 cloned into the construct as a marker for transduction) directed against CD36 or with an empty vector (EV) control and intravenously (i.v.) transferred into TCRα-deficient mice that were i.n. sensitized and rechallenged with HDM extract (**Figure 2H**). TCRα-deficient mice lack T cells but have B cells including GC B cells that allows for IgE production during HDM allergic airway inflammation (Mombaerts et al., 1992). Importantly, we observed significantly reduced CD36 expression in CD4+ T cells in TCRα-deficient mice that received CD36 sgRNA-transduced Cas9-expressing CD4+ T cells compared to TCRα-deficient mice, which have been transferred with EV-transduced CD4+ T cells (**Figure 2I**). Consistent with our previous results using *Cd36^fl/fl^Cd4*-Cre mice, frequencies of mediastinal CD4+ T cells, B cells and GC B cells remained largely unaffected by the transfer of CD36-deleted CD4+ T cells from *Cas9EGFP^stop/fl^Cd4Cre* into TCRα-deficient mice. However, frequencies and the total number of Tfh cells was significantly reduced in mediastinal lymph nodes of mice that underwent HDM allergic airway inflammation. (**Figure 2J-M**). In summary, CD36 is dispensable for germinal center B cell responses and IgE production during HDM-induced airway inflammation, but contributes to sustaining the Tfh cell compartment that gives rise to the pool of terminal differentiated effector T cells in the lung.

### CD36 controls tissue-resident memory T cells during HDM allergic airway inflammation

CD36 was negligible for the Tfh cell-mediated IgE production through GC B cells in mediastinal lymph nodes following HDM airway inflammation. In order to analyze whether CD36 would be important for effector lung T cells we analyzed the lungs of WT and *Cd36^fl/fl^Cd4*-Cre mice that were sensitized and rechallenged with HDM extract. Cell counts and frequencies of lung CD4+ and CD8+ T cell populations and cytokine production appeared unaltered between WT and *Cd36^fl/fl^Cd4*-Cre mice treated with HDM (**Suppl. Figure 10**). However, only in WT mice HDM treatment did trigger a robust expansion of CD44+CD62L-effector CD4+ T cells, whereas this increase was absent in *Cd36^fl/fl^Cd4*-Cre mice (**Figure 2N**). *Ex vivo* bulk transcriptome analysis of total lung CD4+ T cells from HDM treated WT and *Cd36^fl/fl^Cd4*-Cre mice further revealed a significant downregulation of genes related to asthma including *Mx1* and *Gata3* in the absence of CD36 (**Figure 2O**). Gene set enrichment analysis (GSEA) further linked these transcriptional changes to the IL-2–STAT5 pathway (MM3902) and the reactive oxygen species pathway (MM3895), both of which are central drivers of effector differentiation (**Figure 2P**). In addition, these results were of important significance as we have recently reported a strong dependence of Th2 cells on IL-2 signals for their terminal effector differentiation during HDM allergic airway inflammation (Y.-H. Wang et al., 2022). Additionally, analysis of chronically HDM treated WT and *Cd36^fl/fl^Cd4*-Cre mice revealed significant reduced expression of the Th2 cytokine genes *Il4*, *Il5* and *Il13* in lung CD36-deficient CD4+ T cells (**Suppl. Figure 11**). In addition, while most lung immune cell populations remained comparable between *Cd36^fl/fl^Cd4*-Cre mice and WT littermates, CD4+CD69+CD103+ Trm cells represented the only subset that showed a significant reduction in *Cd36^fl/fl^Cd4*-Cre mice, which underwent HDM airway inflammation. (**Figure 2Q**). A defining hallmark of lung CD4+ Trms compared to conventional CD4+ T cells was their elevated expression of the transcription factors PPARγ and GATA3 (**Figure 2R**). Importantly, the expression of both transcription factors, PPARLJ and GATA3, was significantly reduced in CD36-deficient compared to WT Trms (**Figure 2S**). Importantly, while CD36-deficient lung CD4+ Trms showed a significant reduced expression of PPARγ and GATA3 the expression of other Trm transcription factors including Tbet, Blimp1, Notch1, AMPK and mTOR remained unaltered between WT and CD36-deficient Trms. Together, these results demonstrate that CD36 controls the maintenance of lung Trms by sustaining PPARγ and GATA3 expression, thereby linking fatty acid metabolism to long-term effector residency in allergic airway inflammation.

### CD36 deficiency in CD4+ Trms alters gene expression and localization of type-2 conventional dendritic cells during HDM allergic airway inflammation

We found a strong alteration of the frequency and function of CD4+ Trms in *CD36^fl/fl^Cd4*-Cre compared to WT littermate mice during HDM allergic airway inflammation. Trms are known to interact with different adaptive and innate immune cells in the allergic lung. Among these, type-2 conventional dendritic cells (cDC2s) are particularly critical, as they establish direct interactions with Trms and have been shown to promote allergic airway inflammation (Hammad & Lambrecht, 2021; Wu et al., 2024). Using scRNAseq analysis of total CD45+lung cells, we did not observe major changes in the overall composition of innate immune cell populations between WT and *CD36^fl/fl^Cd4*-Cre mice during HDM airway inflammation (**Figure 3A** and **3B**). In contrast, cDC2s displayed pronounced transcriptional alterations in *CD36^fl/fl^Cd4*-Cre mice compared to WT controls (**Figure 3C**). Gene ontology analysis of the DEGs in cDCs2 revealed that the genes including *Cxcl10*, *Il1rn*, *Nfatc2*, *Bsg*, and *Cadm1* belonged to leukocyte migration and oxidative stress (**Figure 3D** and **3E**). For example, *Cadm1* (Cell Adhesion Molecule 1; also known as TSLC1 or Necl-2), an immunoglobulin superfamily adhesion molecule essential for maintaining epithelial integrity and coordinating immune cell–fibroblast interactions, was markedly downregulated in cDC2s from *CD36^fl/fl^Cd4*-Cre mice (Moiseeva et al., 2013). These findings suggested that a deletion of CD36 in CD4+ T cells influences the transcriptional program of cDCs2, potentially modulating their function, e.g. migration and localization, in the allergic lung. To test for this hypothesis, we performed tissue clearing of lungs from WT and *CD36^fl/fl^Cd4*-Cre mice that underwent HDM allergic airway inflammation (**Figure 3F**). 3D reconstructions of intact lungs from WT and *CD36^fl/fl^Cd4*-Cre mice showed increased immune cell infiltration around the airways in WT animals. Immune cell–dense regions were identified based on MHC-II and Sirpα signals, which are characteristic markers enriched in cDC2 populations. In WT mice, cell clustering appeared more frequently and cells formed a visibly thicker layer around inflamed airways compared to the more moderate and patchy infiltration seen in *CD36^fl/fl^Cd4*-Cre lungs (**Figure 3G, I**). Quantitative analysis using a machine-learning-based segmentation confirmed this pattern, showing a higher total volume of inflamed airway regions in WT lungs (**Figure 4H, J**). To account for regional heterogeneity within individual lungs, each lung was subdivided into left and right lobes, and infiltration volumes were determined separately for each lobe. Taken together, these results identify CD36 as a T cell–intrinsic regulator that indirectly shapes the transcriptional program and spatial localization of cDC2s, thereby controlling the extent and architecture of peribronchial immune infiltration during allergic airway inflammation.

**Figure 3:**
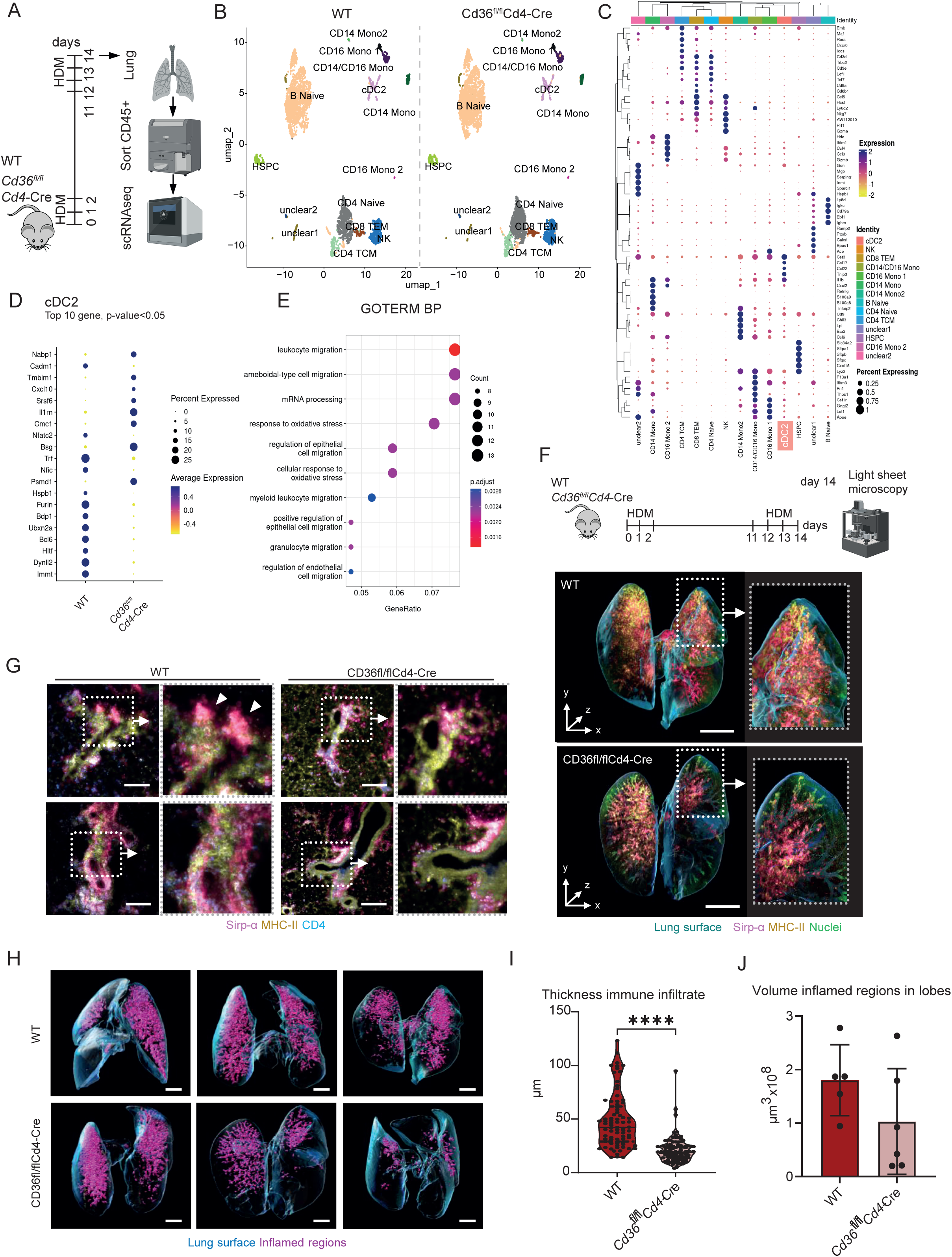
CD4 specific deletion of CD36 leads to DEG in conventional dendritic cell type 2 (cDC2) in HDM induced allergic airway inflammation. Single CD45+ lungs cells of three WT versus three *Cd36^fl/fl^Cd4*-Cre mice upon induction of acute HDM allergic airway inflammation. (A) Depiction of schematic model of acute house dust mite (HDM) allergic airway inflammation followed by CD45+ lungs cells sorting for scRNAseq. (B) Uniform Manifold Approximation and Projection (UMAP) visualization of single CD45+ sputum cells from WT and *Cd36^fl/fl^Cd4*-Cre mice. (C) Dot plot showing the top five marker of each cluster. (D) Dot plot of the top 20 DEGs (differentially expressed genes) having highest logFC and pValue < 0.05 in cDC2s (E) GOTERM BP analysis of DEGs. (F) 3D reconstructions of intact lungs (lung surface, blue) from a wildtype (WT, left) and a CD36fl/flCd4-Cre (KO, right) mouse immunostained for macrophages (Sirp-α, magenta; MHC-II, yellow) and nuclei (green). Scale bar: 2 mm (G) 2D optical slices and zoom-in images showing eribronchial immune cell infiltration by macrophages (Sirp-α, magenta; MHC-II, yellow) and CD4+ T Cells (CD4, cyan) in WT and KO. Scale bar: 100 µm (H) 3D surface rendering (blue) with segmented inflamed airway volume (magenta) in WT and KO lungs based on immunostaining for Sirp-α, MHC-II and CD4. Scale bar: 1.5 mm (I) Quantification of the thickness of the immune cell infiltrate surrounding the airways in WT and KO lungs. Data points represent pooled measurements from nLJ=LJ3 biologically independent samples per group. Black line indicates the median, dotted line the quartiles. Mann Whitney U test. (J) Quantification of inflamed airway volumes per lung lobe. Bars represent mean ± standard deviation; each data point represents a single lung lobe. Two-tailed unpaired t test.

**Figure 4:**
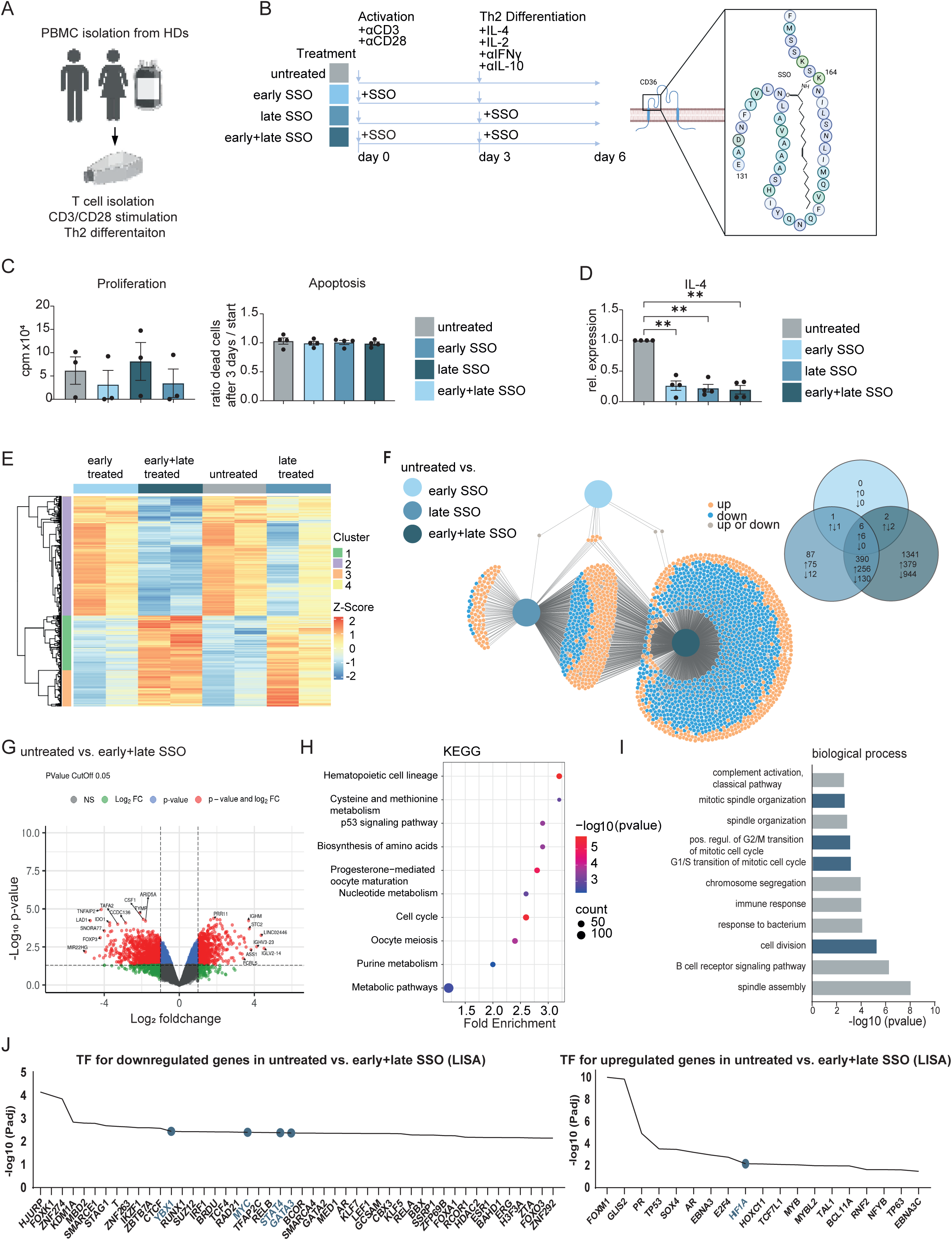
CD36 in human Th2 cells does not critically effect activation and proliferation but it is important during Th2 differentiation via interaction with GATA3, cell cycle and cell division. (A) PBMC isolation from healthy donors for CD4+ T cell isolation. (B) Treatment scheme with sulfosuccinimidyl oleate (SSO) of CD4+ T cells isolated from human peripheral blood mononuclear cells (PBMCs). CD4+ T cells were activated with αCD3 and αCD28 in presence of SSO or absence of SSO After 3 days, activated T cells were cultured in medium containing IL-4, IL-2, αINF_γ_ and αIL-10 for TH2 differentiation in presence or absence of SSO. Readouts were performed on day 6. (C) Development of apoptotic cells on day 6 compared to day 3 and proliferation of T cells during TH2 differentiation. (D) Relative expression of *IL-4* to Treatment A on day 6. (E) Heatmap plot from differentially expressed genes (DEGs) in untreated or SSO treated samples in contrast to all (n=2). Log2FC > 1, FDR < 0.05 (F) DiVenn diagram visualize and compare DEGs lists from treatment B-D with treatment A. Red and blue nodes refer to up and down-regulated DEGs, respectively. Yellow nodes show genes, which are upregulated in one list but downregulated in the compared list. Numbers in black represent the total number of DEGs. Arrows indicates numbers of up or downregulated DEGs. Log2FC > 1. p-value<0.05 (Sun, Liang*, Sufen Dong, Yinbing Ge, Jose P. Fonseca, Zachary Robinson, Kirankumar Mysore, Perdeep Mehta. “DiVenn: An Interactive and Integrated Web-based Visualization Tool for Comparing Gene Lists.” Frontiers in Genetics 10 (2019):421.) (G) Volcano plot depicting DEGs from treatment A compared to D (H) KEGG pathway enrichment analysis of DEGs from treatment A compared to D Log2FC > 1. p-value<0.05 (I) Gene ontology (GO) Biology Function pathways of DEGs from treatment A compared to D based on p-value. Shown is the –log10 of the p-values of the individual pathways. (J) Upstream regulator anaylsis using Landscape in silico analysis (Lisa; http://lisa.cistrome.org/) for downregulated and upregulated genes of DEGs from treatment A compared to D. Sample sizes: (B)-(D) n = 3-4; (E-I) n = 2. Statistical analysis was performed using ordinary one-way ANOVA followed by Tukey’s test for multiple comparisons; *p < 0.05, **p < 0.01, ***p < 0.001.

### CD36 controls the differentiation of human Th2 cells *in vitro*

Our data showed that CD36 is a metabolic checkpoint for T cell tissue residency during a preclinical mouse model of allergic airway inflammation. However, whether CD36 also affects the activation or Th2 differentiation of human T cells remained enigmatic. Therefore, we isolated CD4+ T cells from human PBMCs of unrelated HDs and activated them with activating anti-CD3 and anti-CD28 antibodies in the absence or presence of the CD36 inhibitor sulfosuccinimidyl oleate sodium (SSO) (**Figure 4A** and **4B**). SSO binds irreversibly on CD36 resulting in inhibition of both, signal transduction and fatty acid uptake (Kuda et al., 2013). After 3 days, activated T cells were cultured in medium containing IL-4, IL-2 and anti-IFNγ and αIL-10 antibodies for Th2 differentiation in the absence or presence of SSO. While proliferation and apoptosis were unaffected across all treatment groups (**Figure 4C**), we observed a significant reduction in IL-4 expression in cultures exposed to SSO (**Figure 4D**). To understand this observation in more detail we analyzed the transcriptome of the individual cultures using bulk RNA sequencing (**Figure 4E**). DiVenn analysis of significant DEGs (Log2FC > 1) revealed only 6 genes commonly upregulated across early, late, and combined (early plus late) SSO treatment (**Figure 4F**). Remarkably, early SSO exposure resulted in only 9 DEGs in total, indicating that CD36 inhibition minimally impacts initial T cell activation. In contrast, we detected ∼390 genes to be up- and downregulated in T cells with late SSO treatment during Th2 cell differentiation only and in cells treated early plus late with SSO, during activation and Th2 differentiation, compared to untreated T cells cultures. The strongest transcriptional divergence was observed in untreated versus early plus late SSO conditions, revealing the most number of significant DEGs (**Figure 4G**). KEGG pathway analysis of the latter DEGs revealed altered metabolic and cell cycle pathways upon early and late CD36 inhibition (**Figure 4H**). Additionally, gene ontology highlighted biological processes related to cell division to be significantly affected (**Figure 5I**). In order to identify the key upstream transcription factors regulated CD36, we performed epigenetic Landscape In Silico deletion Analysis (Lisa) (Qin et al., 2020). Using Lisa, we detected several transcription factors important for cell growth and T helper 2 development like YBX1, GATA3, STAT4 and the metabolic regulator Myc connected to DEGs which are downregulated in cells upon CD36 inhibition (**Figure 4J**). In upregulated DEGs of untreated versus eraly plus late SSO, we observe HIF1α to be regulated upon CD36 inhibition. HIF1alpha can regulate several key processes like metabolic reprogramming, cell survival and apoptosis, and cytokine production (**Figure 5J**). Together, these findings establish CD36 as a regulator of human Th2 differentiation, acting primarily during lineage commitment rather than initial T cell activation.

## Discussion

Asthma is a complex and heterogeneous respiratory disease with increasing incidence (Lambrecht & Hammad, 2015). Recent studies have highlighted the critical role of metabolic pathways in regulating immune cell function and allergic airway inflammation. Specifically, metabolic alterations have been associated with the development and maintenance of asthma. For instance, Th2 cells were demonstrated to show a high accessibility of genes related to fatty acid metabolism in mice (Tibbitt et al., 2019). Here, we extend these findings by showing for the first time that asthma patients exhale increased amounts of lipophilic volatile compounds and display altered expression of fatty acid–metabolism–related genes in sputum-derived Th2 cells. Further, our results are in line with studies reporting an increase abundance of lipids in the serum and BAL of patients suffering from allergic asthma (Kang et al., 2014; Ravi et al., 2021; Vasseur & Guillaumond, 2022). Lipids play a crucial role in regulating lung function and inflammation in asthma. Various cell types, including alveolar macrophages, epithelial cells, and T cells, produce a broad spectrum of lipids such as leukotrienes, prostaglandins, lysophosphatidic acids, and surfactant lipids. Notably, airway epithelial type 2 cells (AEC2) generate lipids with both homeostatic and proinflammatory functions. On the one hand, AEC2-derived surfactant lipids contribute to lung homeostasis and the regulation of immune responses. On the other hand, dysregulation of these lipids—many of which also interact with the scavenger receptor CD36—can lead to maladaptive immune responses (Dodd et al., 2016). These observations raise the intriguing possibility that AEC2-derived surfactant lipids engage CD36 on lung Th2 TRMs to support their effector function and long-term tissue residency during allergic airway inflammation. Interestingly, Der p 1, the principal protease of HDM extract, harbors potential cleavage sites within CD36 (data not shown), suggesting a possible mechanism for modulating CD36 function during allergic inflammation.

As the impact of lipid receptor on the pathogenic potential on Th2-like Trms was unknown (Tibbitt et al., 2019), we investigated the role of CD36 *in vitro* and in HDM allergic airway inflammation using conditional knockout mice with a T cell-specific deletion of CD36. By analyzing the thymus and spleen of mice under steady-state conditions, we confirmed that CD36 does not affect thymic T cell development or the composition of splenic cell populations. This provided the rationale to use *CD36^fl/fl^Cd4*-Cre mice to study CD4+ T cell function during HDM-induced allergic airway inflammation. The inconspicuous thymic and splenic phenotype of *CD36^fl/fl^Cd4*-Cre mice suggests that CD36-mediated lipid metabolism is either dispensable for T cell development or compensated by other proteins, such as fatty acid–binding proteins (FABPs) (Zhang et al., 2022). In addition, we observed that CD36 does not affect the T cell function or metabolic activity of Th1 and Th2 cells *in vitro*. This aligns with previous studies, where differences in CD36-dependent T cell regulation were primarily detected in lipid-enriched environments *in vivo*, including the tumor microenvironment, rather than in isolated cell cultures (H. Wang et al., 2020; S. Xu et al., 2021).

Given this, we investigated the role of CD36 in T cells in an HDM-induced allergic airway inflammation model (Ravi et al., 2021, 2021; Tibbitt et al., 2019). In addition, we investigated whether CD36 influences CD4+ T cells prior to their differentiation into terminal Th2 effector cells. To this end, we analyzed mediastinal lymph nodes, the site of early T cell priming following HDM sensitization. Our analysis revealed unchanged Tfh cell and germinal center B cell responses, with no significant alterations in the organization of B and T cell zones. These findings indicate that CD36 is not required for IgE maturation within lymphoid tissues but may instead play a more critical role in sustaining terminal Th2 cells in the inflamed lung environment. These findings were further supported by analyses of human T cells isolated from PBMCs of HDs. CD4+ T cells were activated with anti-CD3 and anti-CD28 in the presence or absence of the CD36 inhibitor SSO, and subsequently differentiated under Th2-polarizing conditions. While pharmacological inhibition of CD36 did not affect initial T cell activation, it markedly impaired the terminal differentiation of human Th2 cells. Thus, CD36 appears dispensable for early T cell activation and priming, but plays a critical role in later stages of Th2 differentiation, underscoring its importance in regulating Th2 metabolism and effector function within inflammatory environments. (Kuda et al., 2013). In addition to Th2 cells, Th17 cells have been implicated in driving therapy-resistant and corticosteroid-insensitive allergic airway inflammation in humans. In our experiments, however, Th17 cell function in the absence of CD36 remained intact both *in vitro* and in the HDM-induced asthma model *in vivo*, and was associated with other and distinct downstream signaling pathways when compared with Th2-like TRMs (data not shown).

In allergic airway inflammation, Trms with a Th2-like phenotype have been shown to play a critical role in sustaining chronic lung inflammation by promoting a type 2 inflammatory environment (Camacho et al., 2022; Rahimi et al., 2020). We found that Trms selectively upregulated PPARLJ and GATA3 and that *Cd36^fl/fl^Cd4*-Cre mice have reduced frequencies of Trms during HDM airway inflammation. Mechanistically, CD36-mediated lipid uptake enhances the expression of the nuclear receptor PPARLJ, promoting the transcription factor GATA3, which is critical for the effector function of Th2 Trm cells in asthma (Stark et al., 2021; S. Xu et al., 2021). This regulatory axis likely contributes to the gene expression of Trm-associated markers such as CD69 and CD103, supports Trm persistence and regulates interaction with type-2 conventional dendritic cells (Jiménez-Fernández et al., 2023; Pan et al., 2017). In this context, PPARLJ binds to the promoter region of CD103 (ITGAE) (data not shown), a hallmark marker of Trms, and cDC2s play an essential role in inducing a Th2-type immune response and were shown to increase in cDC2s in alveolar lavage fluid in a model of bronchial asthma (J. Xu et al., 2024).

Of note, Patel et al. recently demonstrated the importance of CD36 as a critical receptor for phosphorylcholine moieties, underscoring its role in non-lymphoid tissues during asthma (Patel & Kearney, 2017). In line with this, our data support a key contribution of CD36 to shaping local immune responses in the lung, particularly through its impact on tissue-resident T cells. Interestingly, a missense variant in the CD36 gene, identified exclusively in East Asian populations, confers protection against asthma by enhancing CD36 expression in lymphocytes and altering immune protein profiles (Zhi et al., 2025). Together, these findings suggest that CD36 functions as a metabolic checkpoint that establishes and sustains tissue-resident memory Th2 cells during allergic asthma, highlighting its potential as a therapeutic target.

## Limitations of the study

Some limitations should be considered when interpreting our findings. First, while our mouse model of HDM-induced allergic airway inflammation captures key features of human asthma, it may not fully reflect the complexity and heterogeneity of the disease that is characterized by different endotypes in patients. Second, our study primarily relied on T cell–specific CD36 deletion and pharmacological inhibition; thus, compensatory pathways, such as fatty acid– binding proteins (FABPs), may obscure the broader contribution of lipid metabolism to T cell function. Third, our analyses of human cells were restricted to PBMC-derived CD4+ T cells under Th2-polarizing conditions, which may not fully recapitulate the microenvironmental cues present in the inflamed lung, particularly the influence of airway epithelial–derived lipids or dendritic cell interactions. Fourth, while our data suggest that CD36 shapes tissue-resident Th2 cell function via PPARγ and GATA3, the precise lipid species and signaling cascades involved remain to be defined. Together, these limitations highlight the need for future work integrating lipidomic profiling, patient-derived airway samples, and genetic studies to fully delineate the role of CD36 and lipid metabolism in sustaining pathogenic Th2 responses in asthma.

## Material and methods

### Murine CD4+ T cell isolation and culture

Mice were euthanized using CO₂ overdose and cervical dislocation. Spleens and lymph nodes (inguinal, axillary, cervical, and mesenteric) were excised, mechanically dissociated through a 70µm strainer, and washed with PBS/2% FBS. Erythrocytes were lyzed and CD4+ T cells were isolated via negative magnetic selection. Culture plates were coated with rabbit-anti-hamster IgG (MP Biomedicals, #SKU0855398) for 2 h at 37°C before cells were cultured in RPMI medium with 10% FCS, 2 mM GlutaMAX (ThermoScientific, #35050038), 50 µM β-mercaptoethanol (ThermoScientific, # 31350010), 2% penicillin/streptavidin (ThermoScientific, # 15070063) at 37°C, 5% CO₂ in the presence or absence of 50 µg/ml oxLDL. Activation was induced using anti-CD3ε (1 µg/ml) and anti-CD28 (1 µg/ml) (BioLegend, #100340 and #102116, respectively).

### Measuring CD4+ T cell activation, apoptosis and proliferation

CD4+ T cells were activated and differentiated for 3 days. Activation was assessed by staining with CD4, CD62L, CD25, and CD44 in PBS for 20 min at 37°C. Apoptosis was evaluated using an Annexin V/7AAD staining kit (BioLegend, #640930). To this end, cells were stained with CD4, Annexin V, and 7AAD in Annexin Binding Buffer for 20 min at 37 °C. Annexin V detected phosphatidylserine externalization during early apoptosis, while 7AAD identified late apoptotic/necrotic cells. For proliferation analysis, CD4+ T cells were labeled with 5 µM CFSE (Invitrogen, #C34554) for 5 min at room temperature, washed, and cultured for 2–3 days. The cells were stained for CD4 and the dilution of CFSE per cell was analyzed by flow cytometry.

### In vitro Th1 and Th2 cell differentiation

CD4+ T cells were isolated and cultured as described above. For Th1 and Th2 polarization, hIL-2 (30 units/ml), anti-IL-4 (2 µg/ml) and IL-12 (10 ng/ml) or hIL-2 (30 units/ml), anti-IFN-γ (10 µg/ml), anti-Il-12 (10 µg/ml), IL-4 (100 ng/ml) were additionally added to cell cultures, respectively.

### sgRNA CRISPR/Cas9-mediated gene knockout in primary murine CD4+ T cells

Single guideRNA design: Single guide RNAs targeting Cd36 were generated using ChopChop (https://chopchop.cbu.uib.no/). Target gene sequence was obtained from the NCBI Gene Databse. Therefore, the appropriate genome (*mus musculus* (mm10/GRCm38)) was selected in ChopChop to ensure sgRNA specificity. sgRNAs were ranked by ChopChop based on predicted off-target effects (MM0-MM3) and efficiency. Additionally, only sgRNAs with a GC content between 40-60% were considered to ensure stable binding. The ranked sgRNAs suggested by ChopChop were manually reviewed according to the above criteria and, consecutively, selected. sgRNAs were synthesized including forward and backward primers as well as the protospacer adjusted motif (PAM) by eurofins and subsequently cloned into CRISPR backbones.

### CRISPR/Cas9-mediated genetic engineering and transduction

Retro-gRNA-eGFP (225514) and MSCV-IRES-Thy1.1 (17442) DEST vectors were purchased from Addgene. The eGFP moiety in the Retro-gRNA-eGFP vector was excised and replaced with the Thy1.1 moiety from the MSCV-IRES-Thy1.1 DEST vector. The resulting vector was termed MSCV-IRES-eGFP->Thy1.1-DEST_gRNA CD36 and used for further genome editing experiments. For these experiments, viruses were produced by transfecting human embryonic kidney 293T cells with the retroviral packaging construct pCL-ECO and retroviral plasmid containing CD36 sgRNA according to standard protocols (10µg retroviral DNA, 5µg packaging virus, 250mM CaCl_2_·H2O added to 1ml 2X HBS (amount for two cell culture dishes)). DMEM:Chloroquine solution (Chloroquine 1:1000) was prepared and added to the cells, together with 1ml DNA mix as described above. After 7-10h incubation at 37°C, 5% CO2, the DMEM:Chloroquine solution was removed from cells and replaced with 6ml fresh DMEM per 10cm cell culture dish. DMEM was replaced with 6ml fresh DMEM the next day and viral supernatants harvested 24h after the last medium change. CD4+ T cells were isolation from CD4CrexCas9 (C57BL/6N-Gt(ROSA)^26Sorem1(CAG-cas9*,-EGFP)Rsky^/J (JAX stock No: 028551) crossed to Tg(Cd4-cre)1Cwi/BfluJ (JAX stock No: 017336) mice and activated using anti-CD3/CD28 antibodies and Rabbit Anti-Syrian Hamster IgG (H+L) (Jackson ImmunoResearch; #307-005-003) in PBS coated wells. The next day, cells were transduced with previously produced sgRNA-containing viral supernatants by spin infection (2500 rpm, 90 minutes, 32°C) in transduction medium (full RPMI +20ng/ml IL2, +8µg/ml polybrene). Transduced cells received fresh medium (full RPMI +20ng/ml IL2, +2.5ng/ml IL7) the following day and were cultivated an additional 24h at 37°C, 5% CO2. On the fourth day, transduced cells were harvested and analyzed by flow cytometry or used for adoptive T cell transfer into recipient mice.

### Seahorse assays

Mitochondrial respiration and glycolysis were assessed by measuring the oxygen consumption rate (OCR) and extracellular acidification rate (ECAR) using the XFe96 Extracellular Flux Analyzer (Seahorse Bioscience). CD4+ T cells were cultured under Th1 and Th2 conditions for three days, then resuspended in Seahorse XF DMEM (pH 7.4) supplemented with 10 mM glucose, 2 mM glutamine, and 1 mM pyruvate (mitochondrial respiration) or 2 mM glutamine and 1 mM pyruvate (glycolysis). Poly-D-Lysine-coated XFe96 plates were seeded with 200,000 cells per well and centrifuged. OCR was measured following sequential injections of oligomycin (2.5 µM), carbonyl cyanide-4-(trifluoromethoxy)phenylhydrazone (FCCP) (1.5 µM), and rotenone/antimycin (0.5 µM). For glycolysis, ECAR was measured following sequential injections of glucose (2.5 mM), oligomycin (1.5 µM), and 2-deoxy-D-glucose (0.5 mM) for glycolysis. Data were acquired using a XFe96 Extracellular Flux Analyzer (Seahorse Bioscience).

### Induction of house dust mite allergic asthma

Mice were lightly anesthetized with isoflurane and 10 µg house dust mite (HDM) extract in 20 µl PBS was administered intranasally (i.n.). To induce allergic airway inflammation an acute model with i.n. HDM sensitization on the days 0,1, and 3 followed by i.n. HDM rechallenge on days 10–13 and a chronic HDM model with i.n. HDM exposure twice weekly (days 0,1,7,8,14,15,21,22) over four weeks was used. Mice were sacrificed on day 14 (acute) and day 28 (chronic) for analysis, respectively.

### Transfer experiments

Genetic CD36 deletion in murine CD4+ T cells was performed by sgRNA-mediated CRISPR/Cas9 knockout. Two days post-transduction, sgRNA transduced Thy1.1+ live+ CD4+ T cells were sorted and 500,000 cells were intravenously injected into 8–12-week-old TCRα^-/-^(B6.129S2-Tcra^tm1Mom^/J) mice. Subsequently, mice were treated according to the acute house dust mite allergic asthma protocol.

### Tissue collection

Mice were sacrificed by carbon dioxide inhalation and cervical dislocation. The tip of the tail was collected to confirm the genotype by PCR. Mice tissues were processed depending on the subsequent analysis. Blood was collected via cardiac puncture, allowed to clot at room temperature for longer than 30 min and then centrifuged to isolate serum. Bronchoalveolar lavage (BAL) was collected by flushing murine bronchioles with PBS using a tracheal catheter. Retrieved fluid was documented, centrifuged, and supernatants were stored at – 80°C. BAL cells were resuspended in PBS. Mice were perfused with PBS before lung and mediastinal lymph nodes were excised. Lymph node cells were obtained by mechanical dissociation. Upper left lung lobes were fixed in ROTI®Histofix for histopathology analyses. Remaining lung tissue was weighed, digested in Liberase™ for 45 min and further processed as single cell suspension.

### Histopathologic analysis of lung tissue

Fixed tissue was embedded in paraffin and cut into 2 µm thick slices using a microtom (Leica RM-2145). Lung slices were stained with either hematoxylin and eosin (HE) or Periodic acid– Schiff (PAS) including hematoxylin. Thereafter, 1-3 representative sections per animal were selected and the amount of inflammatory cells surrounding the bronchioles or with stained mucus-producing cells within the bronchioles was determined for HE or PAS staining, respectively. The semi-quantitative score for peribronchial cellular infiltration was determined by the percentage of surrounding cells as follows: 0— 0%, 1— <10%, 2— 10-25%, 3— 25-50% and 4— >50%. Mucus production score was determined according to the percentage of PAS stained cells as follows: 0— 0%, 1— <25%, 2— 25-50%, 3— 50-75%, and 4— <75%. Imaging acquisition was performed with the NanoZoomer S360 (Hamamatsu) and analyzed with the NDP.view 2.Ink software (Hamamatsu).

### Flow cytometry analysis

For surface and viability staining, cells from lung and lymph nodes were washed with PBS/2% FCS and incubated in 100 µl of the desired antibody cocktail for 20 min at room temperature in the dark. Subsequently, cells were washed and resuspended in PBS/2% FCS. For intracellular staining, cells were stimulated with 1 μM ionomycin and 20 nM phorbol-12-myristat-13-acetate (PMA) in the presence of 5 µM brefeldin A for 6 h at 37°C. After surface staining, cells were fixed using the eBioscience™ Intracellular Fixation & Permeabilization Buffer Set (Invitrogen™), incubated for 30 min, washed, and stained with intracellular antibody cocktails in 1× Permeabilization Buffer overnight at 4 °C or for 20 min at room temperature. The Foxp3 Transcription Factor Staining Buffer Set (Invitrogen™) was used for transcription factor and nuclear protein staining. Cells were fixed in a 1:4 dilution of Foxp3 Fixation/Permeabilization concentrate for 30 min, washed, and stained in 1× Permeabilization Buffer. For data acquisition cells were transferred in small FACS tubes using the LSR Fortessa flow cytometer including BD FACSDiva 6.1.3 software (BD Bioscience). Single stains and fluorescence minus one (FMO) controls were measured for gating and compensation purpose. Data were analyzed using FlowJoTM software (version 10, BD Biosciences).

### Multiplex immunoassay

Cytokine and chemokine levels in serum and BAL fluid were measured using LEGENDplex™ flow cytometry-based multiplex immunoassay kits in V-bottom plates. Standard curves were generated by serially diluting the top standard (1:4) in assay buffer. Serum samples were diluted 1:2, while BAL fluid was used undiluted. Samples and standards were incubated with antibody-coated beads, followed by biotinylated detection antibodies and PE-streptavidin for quantification. Data were acquired via flow cytometry and analyzed using LEGENDplex™ Qognit software.

### Enzyme-linked immunosorbent assay (ELISA) analysis

Total serum IgE levels were measured using the IgE Mouse Uncoated ELISA Kit (ThermoFisher Scientific, #88-50460). Serum samples were prediluted 1:25 in Assay Buffer A and processed according to the manufacturer’s protocol. Absorbance was measured using a Safire microplate reader (Tecan), and IgE concentrations were determined using a standard curve.

### Quantitative real time reverse transcription-PCR (qRT-PCR)

RNA was extracted using the Quick-RNA™ Miniprep Kit (Zymo Research) according to the manufacturer’s protocol and quantified using a NanoDrop spectrophotometer. Gene expression analysis was performed using the Luna Universal Probe One-Step RT-qPCR Kit (New England Biolabs, #E3006S). RNA samples were diluted to 1 ng/µl and 1 ng per reaction was used. qRT-PCR was conducted on the qTOWER^3^G (Analytik Jena), and relative gene expression was determined using the ΔΔCt method, normalized to *Gapdh* and *Hprt1* (Livak & Schmittgen, 2001).

### Western Blot

CD4+ T cells were isolated and activated using anti-CD3ε (1 µg/ml) and anti-CD28 (1 µg/ml) (BioLegend, #100340 and #102116, respectively) in presence or absence of Dipalmitoylphosphatidylcholin (DPPC) or palmitate for 48 hours.

T cells were lysed in 1% lauryl maltoside, 1% NP-40, 1 mM phenylmethylsulphonyl fluoride, 1LJmM Na3VO4, 10 mM NaF, 10 mM EDTA, 50 mM Tris–HCl (pH 7.5), and 165 mM NaCl.Samples were assayed by sodium dodecyl sulfateLJpolyacrylamide (SDS-PAGE). Proteins were transferred using a semi-dry method onto a nitrocellulose membrane (Amersham Protran, USA). Membranes were blocked in 5% milk and incubated with primary antibodies in 5% bovine serum albumin (BSA) in tris-buffered saline (TBS)/0.2% Tween 20 for 1 h at room temperature (RT) or overnight at 4 °C. Blots were probed with antibodies listed in **Table 2**. Secondary antibodies coupled with horseradish peroxidase (Dianova/Jackson ImmunoResearch Laboratories, Inc., USA) were diluted in 5% milk and incubated for 1 h at RT. Target proteins were detected by ECL (Amersham/GE Healthcare; Chicago, USA). The data were processed and analyzed using the ImageJ software (freeware, http://rsb.info.nih.gov). The total median values of densitometric analysis were used for quantifications.

### Whole mount LN staining and microscopy

Mice were euthanized, and inguinal lymph nodes (LNs) were harvested. The LNs were fixed overnight at 4°C in a 1% PFA fixation buffer (composed of 0.7% w/v Na2HPO4, 1.5% w/v Lysine, 0.2% w/v NaIO4, and 1% PFA in ddH2O). Following this, the LNs were incubated for 5 days in a blocking buffer (0.02% goat serum, 1% BSA, 0.3% Triton X-100, and 0.5 µg/ml Streptavidin in PBS), and then subjected to a 7-day primary antibody staining in the blocking buffer, all at room temperature. T cell zones were stained with a rat monoclonal anti-mouse CD4 (GK1.5) Alexa Fluor 488 conjugate (eBioscience, # 53-0041-82), B cell zones with a rat monoclonal anti-mouse B220 (RA3-6B2) Alexa Fluor 647 conjugate (BD Pharmingen, #557683), and lymphatic structures with a rat monoclonal anti-mouse Lyve1 (ALY7) Biotin conjugate (eBioscience, # 13-0443-82). After antibody staining, the LNs were incubated for 1 day with streptavidin BV421. The LNs were then washed for one day in a wash buffer (0.5% Thioglycerol, 0.2% Triton X-100 in PBS) at room temperature, followed by dehydration in a series of ethanol solutions: 50% ethanol for 2 hours, 70% ethanol overnight, and absolute ethanol for 2 hours. The LNs were then cleared in 100% ethyl cinnamate by shaking at room temperature for 15 minutes and stored in 100% ethyl cinnamate until ready for microscopy. Whole LNs were analyzed using two-photon laser-scanning microscopy while submerged in ethyl cinnamate. The microscopy was performed with a Zeiss 710 upright microscope, using simultaneous detection through four external spectral non-descanned detectors (NDD-Dive module). Illumination was sequentially at 790 and 820nm with a two-photon laser. Bidirectional tile scans (with 25% overlap) of the lymph node were captured as Z stacks (Δz = 5µm), covering the entire LN with an x-y resolution of 1024 x 1024 pixels per image. The raw data were reconstructed, analyzed, and quantified using Imaris (Oxford Instruments, Zurich, Switzerland). T cell zones, B cell zones, and lymphatic tissues were reconstructed using the surface tool, and the volume of the various LN compartments was calculated from the surfaces.

### Human PBMC and CD4+ T cell isolation, activation and measurement of proliferation in the absence and presence of the CD36 inhibitor SSO

Peripheral blood mononuclear cells (PBMCs) were isolated from heparinized blood of healthy donors via Ficoll gradient centrifugation (L6115, Biochrom). Cells were washed in serum-free AIM-V medium (Life Technologies, #12055091) and CD4+ T cells were enriched using a human CD4^+^ T Cell Isolation Kit (Miltenyi, #130-096-533) and an autoMACS® Pro Separator (Miltenyi). 96-well culture plates were pre-coated with scavenger goat anti-mouse IgG (Jackson ImmunoResearch, #115-005-068) overnight at 4°C, washed, and incubated with αCD3e (OKT3) and αCD28 (248.23.2) for 2 hours at 37°C. Purified cells were resuspended in AIM-V medium (1×10LJ cells/ml) cultured in presence or absence of Sulfo-N-succinimidyl Oleate sodium (SSO) (Sigma Aldrich, #SML2148-25MG) and monitored for 3 days using the Incucyte S3® live-cell analysis system (Sartorius). Proliferation was quantified by analyzing cell clusters >1200 µm². Apoptosis was assessed using Incucyte® Cytotox and Caspase-3/7 dyes (Satorius, #4440). On day 3, cells were pulsed with 0.2 µCi/ml [³H]-thymidine (Revvity, #NET027W001MC) for 6 hours, harvested, and [³H]-thymidine incorporation was quantified using a Wallac MicroBeta TriLux scintillation counter (Perkin Elmer).

### Human Th2 differentiation

Proliferating CD4+ T cells were resuspended in fresh AIM-V medium (10LJ cells/ml) and cultured with IL-4 (20 ng/ml), IL-2 (20 ng/ml, PeproTech), anti-IFNγ (2.5 µg/ml), and anti-IL-10 (2.5 µg/ml, BioLegend) in the presence or absence of SSO for 3 days. Cells were either frozen at –80°C or used immediately for further assays. Th2 differentiation was confirmed by IL-5 quantification Quantikine human IL-5 ELISA (bio-techneR&D Systems, #D5000B).

### Sample preparation for bulk RNA sequencing

Samples were generated from CD4+ T cells FACS-sorted from the lungs of WT and *Cd36^fl/fl^Cd4*-Cre mice after acute and chronic HDM allergic airway inflammation treatment or from human Th2 cells cultured in presence or absence of SSO. The cells were sorted for live CD45+CD4+ cells and collected in cooled 1.5 ml tubes containing 200 µl PBS with 10% FBS. RNA was isolated using the RNeasy Plus Mini Kit (QIAGEN, #74136) according to manufacturer’s protocol. The RNA content was measured by using Nano Drop spectrophotometer. RNA samples were sequenced and analyzed in cooperation with Robert Geffers, HZI Braunschweig.

### RNA Sequencing and data Analysis

RNA quality and integrity were assessed using the Agilent 2100 Bioanalyzer (Agilent Technologies, Waldbronn, Germany). Following the manufacturer’s instructions, the NEB SC/Low Input RNA Library Prep Kit (New England BioLabs) was used to create the RNA sequencing library from 1–5 ng of total RNA. The RNA sequencing library was generated from 300ng total RNA using NEBNext® UltraTM II Directional RNA Library Prep Kit for Illumina® (New England BioLabs) and Dynabeads mRNA Direct Micro Kit (Invitrogen) according to manufacturés protocols. The libraries were sequenced on an Illumina NovaSeq 6000 using the NovaSeq 6000 S1 Reagent Kit (100 cycles, paired-end run), generating an average of 5 × 10LJ reads per sample. Each sequence in the raw FASTQ files was trimmed based on base call quality and to remove sequencing adapter contamination. Additionally, sequences shorter than 15 bp were removed using fastq-mcf.

The trimmed reads were then aligned to the reference genome using the open-source short-read aligner STAR (https://code.google.com/p/rna-star/), with parameters specified in the log file. Gene-level counts for each feature in each library were obtained using the Rsubread package (version 2.16.1) in R (version 4.3.3). Gene annotation within the raw count table was performed using the biomaRt package (version 2.58.2). For data preprocessing, only counts with uniquely assigned reads were retained, while those corresponding to rRNA and pseudogenes were excluded. Genes with at least 3 counts per million (CPM) in at least one set of sample replicates were retained. Additionally, libraries with a size deviation exceeding three standard deviations were excluded. The libraries were sequenced on Illumina NovaSeq 6000 using NovaSeq 6000 S1 Reagent Kit (100 cycles, paired end run) with an average of 5 x107 reads per sample Expression data were log₂-transformed and normalized using the trimmed mean of M-values (TMM) method, implemented in the edgeR package (version 4.0.16). Differential gene expression analysis was performed using edgeR. Pathway enrichment analysis (KEGG, GOTERM) was performed using DAVID (https://davidbioinformatics.nih.gov/tools.jsp) (Da Huang et al., 2009; Sherman et al., 2022) or SRPLOT (Tang et al., 2023) with input of differentially expressed genes (DEGs) with official gene IDs. To identify transcriptional regulator predictions, LISA analysis (Qin et al., 2020) was conducted.

### Human sputum collection

Patients were recruited by the Department of Pneumology, University Hospital Magdeburg, Health Campus Immunology, Infectiology and Inflammation, Otto-von-Guericke University Magdeburg, based on predefined inclusion and exclusion criteria. Serum was collected from HDs and asthma patients for total IgE determination. Allergy testing was performed before study inclusion by specific IgE determination according to clinical indications to identify relevant allergens. On the initial study day, induced sputum was obtained after inhalation of hypertonic sodium chloride. Sputum plugs were isolated, treated with a fourfold volume of 0.1% DTT in PBS, homogenized, filtered (70 µm), and centrifuged. Samples were then frozen at −80 °C in IMDM with 10% DMSO and 40% FBS. The following day, the first breath sample (t_bp_) was collected using a 3 Liter Tedlar bag. Airway hyperresponsiveness was assessed via non-specific methacholine provocation test. Subsequently, specific allergen provocation was performed using ascending concentrations of sodium chloride solution or the identified allergen under resting breathing conditions in HDs or asthma patients, respectively. Lung function was measured within 10 min following each inhalation. Allergen challenge was terminated if forced expiratory volume in 1 second (FEV₁) decreased by ≥20% from baseline or airway resistance increased by ≥100%, indicating a positive response. A second breath sample was obtained immediately post-provocation (t_0_), and a third sample was collected 8 h (t_8_) after sodium chloride solution or allergen exposure.

### Breath sample collection and VOC Analysis

Breath samples were collected in 3 Liter Tedlar bags. Participants were carefully instructed to avoid consumption of odorous drinks and food prior to breath sampling to minimize the influence of external VOCs on the concentration of exhaled compounds. The samples were analyzed within 2-4 hours after collection at the Institute of Medical Technology to prevent the loss of gas compounds from the sampling bags. Breath gas analysis was performed using a commercial PTR-MS with a Time-of-Flight (TOF) mass detector (PTR-TOF 2000, Ionicon Analytik, Innsbruck, Austria), which offers sensitive offline and online measurements in the low ppb range. The breath sampling followed the procedures outlined by Gbaoui et al. 2022 (Gbaoui et al., 2022). PTR-MS measurements were carried out with a drift tube pressure of 2.3 mbar and VOC masses were analyzed in consecutive scans across a mass-to-charge ratio range from m/z 20 to 200. To account for background VOCs, ambient air samples were collected from each sampling room. Additionally, salbutamol-containing samples were analyzed to assess potential residual drug presence in exhaled air.

### Sample preparation and single cell RNA sequencing

Sputum samples from HDs and asthma patients (n=3) were incubated with FC-block (Biolegend, #422301) and labeled with unique TotalSeq-C hashtag antibodies (C0251-C0256 anti-human, 1:100, BioLegend). Samples were fluorescently labeled with AnnexinV-APC and CD45-FITC (Biolegend, #640930, #304006) and sorted for live CD45+ cells using an Aria III fluorescence-activated cell sorter (BD). Single cell suspension of lungs collected from WT versus *Cd36^fl/fl^Cd4*-Cre (n=3) mice upon induction of acute HDM allergic airway inflammation were incubated with FC-block (Biolegend, #422301) before labeling with unique TotalSeq-C hashtag antibodies (C0301-C0306 anti-mouse, 1:100, BioLegend). Fluorescent-labeled samples were sorted for negative AnnexinV-APC and positive CD3-BV421, CD8-BV421, CD11b-BV421, CD11c-BV421, F4/80-BV421, MHCII-BV421, B220-BV421, CD4-PB signal. The Chromium Next GEM Single Cell 5’ Reagent Kits v2 (Dual Index) workflow was employed for single-cell gene expression analysis. Cell suspensions were loaded onto a Chromium Next GEM Chip, containing Gel Beads and Master Mix, to encapsulate individual cells into nanoliter-scale Gel Bead-in-Emulsions (GEMs). Reverse transcription was performed to generate barcoded cDNA from polyadenylated RNA. Hashtag antibodies were utilized for sample multiplexing. Cells were stained with TotalSeq-C hashtag antibodies, washed, and processed through the Chromium workflow, which captured antibody tags along with cellular mRNA. Feature Barcode libraries were selectively amplified, and barcoded cDNA was subsequently amplified via PCR. Libraries were constructed according to 10x Genomics protocols, incorporating dual-indexed adapters for multiplexed sequencing. Separate libraries were prepared for gene expression and Feature Barcode data. Sequencing was conducted on an Illumina NovaSeq6000 (100 cycles, asynchronous) with recommended read depths of approximately 50,000 reads per cell for gene expression and 5,000 reads per cell for Feature Barcode libraries. The targeted cell number for encapsulation was 10,000 cells.

### Single Cell RNA sequencing data analysis (human sputum)

The obtained scRNAseq data using the 10X Genomics Chromium technology were quantified and reads aligned using the Cell Ranger Pipeline provided by 10x genomics. The resulting gene count matrices (UMIs per cell and gene) were further processed using R package Seurat version 5.3.0 (Hao et al., 2024). A Seurat Object was created requiring a gene to be expressed in a minimum of 3 cells and a cell having a minimum of 100 features (16,065 features, 1,810 cells). To filter empty droplets, doublets and dead cells, we apply further filtration. Cells get excluded if (1) the number of unique genes per cell nFeature_RNA≤100 or nFeature_RNA≥3000; (2) the total number of molecules within a cell nCount_RNA≤700 or nCount_RNA≥15000; (3) the percentage of reads mapping to mitochondrial genes percent mt≥10 resulting in a total of 1,233 cells. The HTODemux() function was employed for demultiplexing samples labeled with Hashtag oligonucleotides (HTOs). A new HTO assay was added, and data normalized using centered log ratio transformation (CLR). Subsequently, demultiplexing was performed with default configuration (positive.quantile=0.99). Downstream analyses were only performed for those cells identified as singlets by the algorithm. Data were normalized (method “LogNormalize”), the 2,000 most variable features were determined (method “vst”) and data scaled according to these features. For visualization and clustering, Principal Component Analysis (PCA) was used to generate uniform manifold approximation and projection (UMAP) coordinates (dimensions 1 to 10, seed=42). Nearest neighbors (dimensions 1 to 10) and clusters (resolution = 0.5) were identified. Celltype annotation was performed manually, based on the presence of marker genes. Annotation was aided by R package SingleR and DatabaseImmuneCellExpressionData, as well as the web application PuMA. To identify subtypes of cells within the T cell cluster, we calculated module scores for feature expression programs using Seurat. We defined the modules CD3CD4+ and CD3CD4GATA3+. Cells were assigned to the highest scoring module >0. In case no cell in the T cell cluster features a score >0, the cell is labeled as “other T cell”.

### Single Cell RNA sequencing data analysis (mouse lung)

Sequencing data were processed using Cell Ranger (version 7.0, 10x Genomics). The resulting gene count matrix, containing unique molecular identifiers (UMIs) per cell and gene, was imported into R and analyzed with the Seurat package (version 4.9.9.9045) using the Read10X function. For sample demultiplexing, cells labeled with hashtag oligonucleotides (HTOs) were processed using the HTODemux function.This function first performs k-medoid clustering on the normalized HTO values, separating cells into K+1 clusters, where K represents the number of samples. For each HTO, a ‘negative’ distribution is computed by identifying the cluster with the lowest average value as the negative group. A negative binomial distribution is then fitted to this negative cluster, and the 0.9 quantile of this distribution is used as a threshold for classification. Cells are classified as positive or negative for each HTO based on these thresholds. Cells positive for more than one HTO are flagged as doublets and excluded from further analysis. Preprocessing of the expression data matrix was performed on a sample-wise basis. Cells with low feature counts (< 500) and high mitochondrial gene content (> 5%) were excluded (leaky cells). Data normalization and integration were performed in R v4.2.1 using Seurat v4.9.9.9045 with the SCTransform(), SelectIntegrationFeatures(), PrepSCTIntegration(), FindIntegrationAnchors(), and IntegrateData() functions. For visualization and clustering, Principal Component Analysis (PCA) was used to generate two-dimensional t-distributed stochastic neighbor embedding (t-SNE) and uniform manifold approximation and projection (UMAP) coordinates, visualized with RunTSNE() and RunUMAP(), respectively. Nearest neighbors and clusters were identified with FindNeighbors() and FindClusters() (clustering resolution = 0.5), and hierarchical clustering was performed with BuildClusterTree(). Cluster annotation was conducted either manually or automatically using Seurat’s Reference-Based Label Transfer/Mapping with cell-type specific marker sets/references. Cluster and cell-type specific marker genes were identified using the FindAllMarkers() function.

### Tissue clearing and whole mount staining of lungs

Whole lungs were fixed with 4% PFA overnight and washed twice with PBS for one hour each at 4°C. The lungs were then decolorized in Quadrol (25% in H2O) for 48 hours at 37°C while shaking to remove the heme and afterwards washed two times for 30 minutes with PBS at 4°C. For enhancement of antibody penetration, samples were incubated in penetration buffer (20% DMSO (Sigma-Aldrich), 0.3 M glycine (Sigma-Aldrich) in PBS with 0.2% Trition-X100 (Sigma-Aldrich)) at 4°C under constant shaking for 24 h. The samples were then bleached with an increasing methanol (MeOH, Sigma-Aldrich) series (30%, 50%, 70% for 2 h and 95%, 100% for 90 min each at 4°C), followed by incubation in bleaching solution (MeOH + 17% DMSO + 5% hydrogen peroxide (H2O2, Carl Roth)) over night at 4°C. Rehydration was performed with the MeOH series in reverse order (100%, 95% for 30 min and 70%, 50%, 30% for 2h each) at 4°C. After the MeOH series, samples were washed three times in washing buffer (0.2% Triton X-100 in PBS) for 30 min each at room temperature. Lungs were then transferred to blocking buffer (10% DMSO, 6% FCS, 0.2% Triton X-100 in PBS) and incubated for 24 h at 37°C with constant shaking. Antibodies were diluted in antibody buffer (5% DMSO, 3% FCS, 0.2% Tween20 (Sigma-Aldrich) and 0.05% sodium azide in PBS) and the samples were incubated at 37°C with constant shaking for 7 days. After the incubation, lungs were washed 6 times with washing buffer (0.2% Tween, 100µg/ml Heparin and 0.05% sodium azide in PBS) at room temperature with an additional washing step over night. The samples were incubated in antibody buffer with secondary antibodies and dyes for 7 days at 37°C in the dark while shaking. The samples were stained with anti-CD4 (clone REA1211, Miltenyi Biotec), anti-Sirpa-AlexaFluor594 (clone P84, BioLegend) and anti-MHC-II-AlexaFluor700 (clone M5/114.15.2, ThermoFisher). After seven washes with the same time intervals and buffer as described above, the lungs were embedded in agarose (1% in distilled H2O, ThermoFisher) and dehydrated at 4 °C in a graded ethanol (EtOH) series (pH>9, 50%, 70%, for 12 h each, 100%, 100%, for 24 h each). For refractive index matching, samples were transferred to ethyl cinnamate (ECi, Sigma-Aldrich) and incubated for 24 h with refreshment of the solution after 12 h. Light-sheet microscopy image acquisition and microscopic image analysis: The samples were recorded at the Blaze Ultramicroscope II (Miltenyi Biotec, Bergisch Gladbach, Germany), using a 1x objective (NA 0.1) and 2.5x magnification. Images were recorded with a 4.2 Megapixel sCMOS camera. The following filter combinations were used for excitation (ex) and emission (em): ex 470/40 and em 525/50 for SYTOX Green (nuclei), ex 595/21 and em 630/30 for Alexa Fluor 594 (SIRPα), ex 630/20 and em 667/30 for Vio R667 (CD4) and ex 650/45 and em 720/60 for Alexa Fluor 700 (MHC II). Light-sheet images were reconstructed using Imaris 10.1 (Bitplane AG).

### Quantification of light-sheet images

Images were stitched, processed and quantified using Imaris 10.1.1 (Bitplane AG). The lung surfaces of scans acquired at low resolution were segmented using the built-in machine-learning based pixel classification tool. Foreground and background pixels were determined by training the model on all channels and surfaces were calculated with a surface grain size of 75 µm. The lung surface was then split into the lung lobes to obtain statistics. Inflamed airways were similarly segmented using the machine-learning tool and training the model on all channels. The model was trained on one WT and one KO sample and was then applied to all other scans. The surface gain size used was 15 µm and surfaces with volumes below 1000 voxels were excluded. We further acquired regions at high resolution and segmented the inflamed regions surrounding airwas based on the high nuclear density of the immune infiltrate using the machine-learning tool. The surface was then manually segmented into 5 discrete sections and quantified regarding signal intensity of MHC-II, CD4, Sirpa and normalized to the volume.

### Normalization

If not otherwise indicated data was normalized to the mean of the untreated wild type control mice.

### Ethics declaration

All procedures were approved by local government agencies to S.K. for the HDM model (Landesverwaltungsamt Sachsen-Anhalt, 42502-2-1633 UniMD). All methods were carried out in accordance with relevant guidelines and regulations and are reported in accordance with ARRIVE guidelines. Patient sample analyses was approved by local ethics committee (study number 123/23).

### Statistical analysis

All data were analyzed using GraphPad Prism 8.0 (GraphPad software, Inc.) and are presented as the meanLJ±LJSEM, if not indicated otherwise. Comparisons of two groups was analyzed by unpaired student’s t test whereas comparisons across multiple groups were performed by ordinary one-way ANOVA. P valuesLJ<LJ0.05 were considered as statistically significant; different levels of significance were indicated as follows: *PLJ<LJ0.05; **PLJ<LJ0.01; ***PLJ<LJ0.001.

## Supporting information

Table 1 characteristics of patients

Table 2 Antibodies and reagents

## Author contributions

Design and principal investigator of the study: S.K.; Drafting the paper: A.K. and S.K.; Writing the paper: A.K. and S.K.; Performing experiments: A.K., S.S., N.J.-N., A.S., J.D., K. K.-D., A.G., O. H., L.B., T.F., J.N., C.M., A.M., M.B., M.F., S. S.-K., B.W., S.K.; Analyzing data: A.K., S.S., N.J.-N., A.S., J.D., K. K.-D., A.G., L.Z., T.F., J.N., A.M., R.G., S. S.-V., M. F., S.S.K., B.W., and S.K.; Critical academic input, infrastructure, discussion and interpretation of the data: B.S., J.S., T.T., D.M., J.V., J.S., A. Mü., D.R., S. S.-K., A.D., and S.K.; All authors reviewed the manuscript and agreed to the submission.

## References

1. Ballesteros-Tato, A., Randall, T. D., Lund, F. E., Spolski, R., Leonard, W. J., & León, B. (2016). T Follicular Helper Cell Plasticity Shapes Pathogenic T Helper 2 Cell-Mediated Immunity to Inhaled House Dust Mite. Immunity, 44(2), 259–273. 10.1016/j.immuni.2015.11.017

2. Beggs, P. J., Clot, B., Sofiev, M., & Johnston, F. H. (2023). Climate change, airborne allergens, and three translational mitigation approaches. EBioMedicine, 93, 104478. 10.1016/j.ebiom.2023.104478

3. Camacho, D. F., Velez, T. E., Hollinger, M. K., Wang, E., Howard, C. L., Darnell, E. P., Kennedy, D. E., Krishack, P. A., Hrusch, C. L., Clark, M. R., Moon, J. J., & Sperling, A. I. (2022). Irf4 expression by lung dendritic cells drives acute but not Trm cell-dependent memory Th2 responses. JCI Insight, 7(21). 10.1172/jci.insight.140384

4. Chen, Y., Zhang, J [Jue], Cui, W., & Silverstein, R. L. (2022). Cd36, a signaling receptor and fatty acid transporter that regulates immune cell metabolism and fate. The Journal of Experimental Medicine, 219(6). 10.1084/jem.20211314

5. Da Huang, W., Sherman, B. T., & Lempicki, R. A. (2009). Systematic and integrative analysis of large gene lists using DAVID bioinformatics resources. Nature Protocols, 4(1), 44– 57. 10.1038/nprot.2008.211

6. Dodd, C. E., Pyle, C. J., Glowinski, R., Rajaram, M. V. S., & Schlesinger, L. S. (2016). Cd36-Mediated Uptake of Surfactant Lipids by Human Macrophages Promotes Intracellular Growth of Mycobacterium tuberculosis. Journal of Immunology (Baltimore, Md. : 1950), 197(12), 4727–4735. 10.4049/jimmunol.1600856

7. Fachet, M., Lowitzki, S., Reckzeh, M.lzlL., Walles, T., & Hoeschen, C. (2022). Investigation of everyday influencing factors on the variability of exhaled breath profiles in healthy subjects. Current Directions in Biomedical Engineering, 8(2), 261–264. 10.1515/cdbme-2022-1067

8. Gbaoui, L., Fachet, M., Lüno, M., Meyer-Lotz, G., Frodl, T., & Hoeschen, C. (2022). Breathomics profiling of metabolic pathways affected by major depression: Possibilities and limitations. Frontiers in Psychiatry, 13, 1061326. 10.3389/fpsyt.2022.1061326

9. Hammad, H., & Lambrecht, B. N [Bart N.] (2021). The basic immunology of asthma. Cell, 184(6), 1469–1485. 10.1016/j.cell.2021.02.016

10. Hao, Y., Stuart, T., Kowalski, M. H., Choudhary, S., Hoffman, P., Hartman, A., Srivastava, A., Molla, G., Madad, S., Fernandez-Granda, C., & Satija, R. (2024). Dictionary learning for integrative, multimodal and scalable single-cell analysis. Nature Biotechnology, 42(2), 293–304. 10.1038/s41587-023-01767-y

11. Höglund, N., Nieminen, P., Mustonen, A.lzlM., Käkelä, R., Tollis, S., Koho, N., Holopainen, M., Ruhanen, H., & Mykkänen, A. (2023). Fatty acid fingerprints in bronchoalveolar lavage fluid and its extracellular vesicles reflect equine asthma severity. Scientific Reports, 13(1), 9821. 10.1038/s41598-023-36697-x

12. Jiménez-Fernández, M., La Fuente, H. de, Martín, P., Cibrián, D., & Sánchez-Madrid, F. (2023). Unraveling CD69 signaling pathways, ligands and laterally associated molecules. 10.17179/EXCLI2023-5751

13. Kang, Y. P., Lee, W. J., Hong, J. Y., Lee, S. B., Park, J. H., Kim, D., Park, S., Park, C.lzlS., Park, S.lzlW., & Kwon, S. W. (2014). Novel approach for analysis of bronchoalveolar lavage fluid (BALF) using HPLC-QTOF-MS-based lipidomics: Lipid levels in asthmatics and corticosteroid-treated asthmatic patients. Journal of Proteome Research, 13(9), 3919–3929. 10.1021/pr5002059

14. Kuda, O., Pietka, T. A., Demianova, Z., Kudova, E., Cvacka, J., Kopecky, J., & Abumrad, N. A. (2013). Sulfo-N-succinimidyl oleate (SSO) inhibits fatty acid uptake and signaling for intracellular calcium via binding CD36 lysine 164: Sso also inhibits oxidized low density lipoprotein uptake by macrophages. The Journal of Biological Chemistry, 288(22), 15547–15555. 10.1074/jbc.M113.473298

15. Lambrecht, B. N [Bart N.], & Hammad, H. (2015). The immunology of asthma. Nature Immunology, 16(1), 45–56. 10.1038/ni.3049

16. Lambrecht, B. N [Bart N.], Hammad, H., & Fahy, J. V. (2019). The Cytokines of Asthma. Immunity, 50(4), 975–991. 10.1016/j.immuni.2019.03.018

17. Livak, K. J., & Schmittgen, T. D. (2001). Analysis of relative gene expression data using real-time quantitative PCR and the 2(-Delta Delta C(T)) Method. *Methods (San Diego*, Calif.), 25(4), 402–408. 10.1006/meth.2001.1262

18. Loisel, D. A., Du, G., Ahluwalia, T. S., Tisler, C. J., Evans, M. D., Myers, R. A., Gangnon, R. E., Kreiner-Møller, E., Bønnelykke, K., Bisgaard, H., Jackson, D. J., Lemanske, R. F., Nicolae, D. L., Gern, J. E., & Ober, C. (2016). Genetic associations with viral respiratory illnesses and asthma control in children. Clinical and Experimental Allergy : Journal of the British Society for Allergy and Clinical Immunology, 46(1), 112–124. 10.1111/cea.12642

19. Ma, X., Xiao, L., Liu, L., Ye, L., Su, P., Bi, E., Wang, Q [Qiang], Yang, M., Qian, J., & Yi, Q. (2021). Cd36-mediated ferroptosis dampens intratumoral CD8+ T cell effector function and impairs their antitumor ability. Cell Metabolism, 33(5), 1001–1012.e5. 10.1016/j.cmet.2021.02.015

20. Moiseeva, E. P., Roach, K. M., Leyland, M. L., & Bradding, P. (2013). Cadm1 is a key receptor mediating human mast cell adhesion to human lung fibroblasts and airway smooth muscle cells. PloS One, 8(4), e61579. 10.1371/journal.pone.0061579

21. Mombaerts, P., Clarke, A. R., Rudnicki, M. A., Iacomini, J., Itohara, S., Lafaille, J. J., Wang, L., Ichikawa, Y., Jaenisch, R., & Hooper, M. L. (1992). Mutations in T-cell antigen receptor genes alpha and beta block thymocyte development at different stages. Nature, 360(6401), 225–231. 10.1038/360225a0

22. Okamura, D. M., Pennathur, S., Pasichnyk, K., López-Guisa, J. M., Collins, S., Febbraio, M., Heinecke, J., & Eddy, A. A. (2009). Cd36 regulates oxidative stress and inflammation in hypercholesterolemic CKD. Journal of the American Society of Nephrology : JASN, 20(3), 495–505. 10.1681/ASN.2008010009

23. Pan, Y., Tian, T., Park, C. O., Lofftus, S. Y., Mei, S., Liu, X., Luo, C., O’Malley, J. T., Gehad, A., Teague, J. E., Divito, S. J., Fuhlbrigge, R., Puigserver, P., Krueger, J. G., Hotamisligil, G. S., Clark, R. A., & Kupper, T. S. (2017). Survival of tissue-resident memory T cells requires exogenous lipid uptake and metabolism. Nature, 543(7644), 252–256. 10.1038/nature21379

24. Patel, P. S., & Kearney, J. F. (2017). Cd36 and Platelet-Activating Factor Receptor Promote House Dust Mite Allergy Development. *Journal of Immunology (Baltimore*, Md*. :* 1950), *199*(3), 1184–1195. 10.4049/jimmunol.1700034

25. Qin, Q., Fan, J., Zheng, R., Wan, C., Mei, S., Wu, Q., Sun, H., Brown, M., Zhang, J [Jing], Meyer, C. A., & Liu, X. S. (2020). Lisa: Inferring transcriptional regulators through integrative modeling of public chromatin accessibility and ChIP-seq data. Genome Biology, 21(1), 32. 10.1186/s13059-020-1934-6

26. Rahimi, R. A., Nepal, K., Cetinbas, M., Sadreyev, R. I., & Luster, A. D. (2020). Distinct functions of tissue-resident and circulating memory Th2 cells in allergic airway disease. The Journal of Experimental Medicine, 217(9). 10.1084/jem.20190865

27. Ravi, A., Goorsenberg, A. W., Dijkhuis, A., Dierdorp, B. S., Dekker, T., van Weeghel, M., Sabogal Piñeros, Y. S., Shah, P. L., Hacken, N. H. ten, Annema, J. T., Sterk, P. J., Vaz, F. M., Bonta, P. I., & Lutter, R. (2021). Metabolic differences between bronchial epithelium from healthy individuals and patients with asthma and the effect of bronchial thermoplasty. Journal of Allergy and Clinical Immunology, 148(5), 1236– 1248. 10.1016/j.jaci.2020.12.653

28. Safran, M., Dalah, I., Alexander, J., Rosen, N., Iny Stein, T., Shmoish, M., Nativ, N., Bahir, I., Doniger, T., Krug, H., Sirota-Madi, A., Olender, T., Golan, Y., Stelzer, G., Harel, A., & Lancet, D. (2010). Genecards Version 3: The human gene integrator. Database : The Journal of Biological Databases and Curation, 2010, baq020. 10.1093/database/baq020

29. Sherman, B. T., Hao, M., Qiu, J., Jiao, X., Baseler, M. W., Lane, H. C., Imamichi, T., & Chang, W. (2022). David: A web server for functional enrichment analysis and functional annotation of gene lists (2021 update). Nucleic Acids Research, 50(W1), W216–W221. 10.1093/nar/gkac194

30. Stark, J. M [Julian M.], Coquet, J. M [Jonathan M.], & Tibbitt, C. A [Christopher A.] (2021). The Role of PPAR-γ in Allergic Disease. Current Allergy and Asthma Reports, 21(11), 45. 10.1007/s11882-021-01022-x

31. Tang, D., Chen, M [Mingjie], Huang, X [Xinhua], Zhang, G [Guicheng], Zeng, L., Zhang, G [Guangsen], Wu, S., & Wang, Y. (2023). Srplot: A free online platform for data visualization and graphing. PloS One, 18(11), e0294236. 10.1371/journal.pone.0294236

32. Tibbitt, C. A [Christopher Andrew], Stark, J. M [Julian Mario], Martens, L., Ma, J., Mold, J. E., Deswarte, K., Oliynyk, G., Feng, X., Lambrecht, B. N [Bart Norbert], Bleser, P. de, Nylén, S., Hammad, H., Arsenian Henriksson, M., Saeys, Y., & Coquet, J. M [Jonathan Marie] (2019). Single-Cell RNA Sequencing of the T Helper Cell Response to House Dust Mites Defines a Distinct Gene Expression Signature in Airway Th2 Cells. Immunity, 51(1), 169–184.e5. 10.1016/j.immuni.2019.05.014

33. Vasseur, S., & Guillaumond, F. (2022). Lipids in cancer: A global view of the contribution of lipid pathways to metastatic formation and treatment resistance. Oncogenesis, 11(1), 46. 10.1038/s41389-022-00420-8

34. Wang, H., Franco, F., Tsui, Y.lzlC., Xie, X., Trefny, M. P., Zappasodi, R., Mohmood, S. R., Fernández-García, J., Tsai, C.lzlH., Schulze, I., Picard, F., Meylan, E., Silverstein, R., Goldberg, I., Fendt, S.lzlM., Wolchok, J. D., Merghoub, T., Jandus, C., Zippelius, A., & Ho, P.lzlC. (2020). Cd36-mediated metabolic adaptation supports regulatory T cell survival and function in tumors. Nature Immunology, 21(3), 298–308. 10.1038/s41590-019-0589-5

35. Wang, Y.lzlH., Noyer, L., Kahlfuss, S., Raphael, D., Tao, A. Y., Kaufmann, U., Zhu, J., Mitchell-Flack, M., Sidhu, I., Zhou, F., Vaeth, M., Thomas, P. G., Saunders, S. P., Stauderman, K., Curotto de Lafaille, M. A., & Feske, S. (2022). Distinct roles of ORAI1 in T cell-mediated allergic airway inflammation and immunity to influenza A virus infection. Science Advances, 8(40), eabn6552. 10.1126/sciadv.abn6552

36. World Health Organization. (2024). https://www.who.int/news-room/fact-sheets/detail/asthma?utm_source=chatgpt.com

37. Wu, C., Yu, H., Liang, F., Huang, X [Xiancong], Jiang, B., Lou, Z., Liu, Y., Wu, Z., Wang, Q [Qi], Shen, H., Chen, M [Ming], Wu, P., & Wu, M. (2024). Hypoxia inhibits the iMo/cDC2/CD8+ TRMs immune axis in the tumor microenvironment of human esophageal cancer. Journal for Immunotherapy of Cancer, 12(7). 10.1136/jitc-2024-008889

38. Xu, J., Cao, S., Xu, Y., Chen, H., Nian, S., Li, L [Lin], Liu, Q., Xu, W., Ye, Y., & Yuan, Q. (2024). The role of DC subgroups in the pathogenesis of asthma. Frontiers in Immunology, 15, 1481989. 10.3389/fimmu.2024.1481989

39. Xu, S., Chaudhary, O., Rodríguez-Morales, P., Sun, X., Chen, D., Zappasodi, R., Xu, Z., Pinto, A. F. M., Williams, A., Schulze, I., Farsakoglu, Y., Varanasi, S. K., Low, J. S.,Tang, W., Wang, H., McDonald, B., Tripple, V., Downes, M., Evans, R. M., . . . Kaech, S. M. (2021). Uptake of oxidized lipids by the scavenger receptor CD36 promotes lipid peroxidation and dysfunction in CD8+ T cells in tumors. Immunity, 54(7), 1561–1577.e7. 10.1016/j.immuni.2021.05.003

40. Zhang, M., Lin, X., Yang, Z., Li, X., Zhou, Z., Love, P. E., Huang, J., & Zhao, B. (2022). Metabolic regulation of T cell development. Frontiers in Immunology, 13, 946119. 10.3389/fimmu.2022.946119

41. Zhi, L., Zheng, Q., Jiang, Y., Yu, L., Li, L [Linzehao], Song, Y., Peng, B., Zhang, C., Jiang, H., Li, R., Mentch, F., Glessner, J., Jia, P., Tang, H., Hakonarson, H., & Chang, X. (2025). Multi-trait genetic analysis of asthma and eosinophils uncovers pleiotropic loci in East Asians. Nature Communications, 16(1), 5081. 10.1038/s41467-025-60405-0

