## Supplementary material for "CD36 is a metabolic checkpoint for Th2 cell tissue residency during allergic airway inflammation": Table 1 characteristics of patients

**Table 1: Characteristics patients**

| **ID** | **Group** | **Sex** | **Age** | **Weight** | **Hight** | **BMI** | **Allergen** | **Total IgE** | **specific IgE** |
| --- | --- | --- | --- | --- | --- | --- | --- | --- | --- |
| A13 | Asthma | w | 39 | 106 | 162 | 40.4 | *Dermatophagoides pteronyssinus* | 165 | 11.4 |
| A16 | Asthma | m | 25 | 85 | 187 | 24.3 | *Dermatophagoides farinae* | 106 | 11.4 |
| A18 | Asthma | w | 38 | 52 | 161 | 21.1 | Timothy grass | 109 | 28.4 |
| K1 | HD | w | 26 | 65 | 157 | 26.4 | - | 6.7 | - |
| K11 | HD | m | 26 | 88 | 178 | 27.8 | - | 11.7 | - |
| K12 | HeHD | w | 22 | 52 | 164 | 19.3 | - | 9.1 | - |
