## Supplementary material for "CD36 is a metabolic checkpoint for Th2 cell tissue residency during allergic airway inflammation": Table 2 Antibodies and reagents

| **Antibody (Western blot)** | **Company** | **Cat. No.** |
| --- | --- | --- |
| Total OXPHOS Rodent WB Antibody Cocktail | Abcam | ab110413 |
| PGC-1a (3G6) Rabbit mAb | Cell Signaling | 2178 |
| Phospho-AMPKa (Thr 172) (40H9) | Cell Signaling | 2535 |
| AMPKa | Cell Signaling | 2793 |
| HIF-1alpha | Cell Signaling | 36169 |
| Lipin1 | Cell Signaling | 5195 |
| mTor | Cell Signaling | 2983 |
| phospho-mTor | Cell Signaling | 2971 |
| NFAT-C1 | Enzo Life Science | 804-022-1R 100 |
| Blimp-1 | Invitrogen | 53-9850-82 |
| PPARY | SIGMA | MA5-14889 |
| B-Actin | SIGMA | A5441-100UL |

| **Antibody**  **(FACS)** | **Clone** | **Fluorochrom** | **Company** | **Cat.No.** |
| --- | --- | --- | --- | --- |
| CD4 | RM4-5 | PB | Biolegend | 100531 |
| CD11b | M1/70 | PB | Biolegend | 101224 |
| CD25 | PC61 | FITC | Biolegend | 102006 |
| CD4 | GK1.5 | FITC | Biolegend | 100406 |
| CD62L | MEL-14 | FITC | Biolegend | 104406 |
| GL7 | GL7 | FITC | Biolegend | 144603 |
| CD69 | H1.2F3 | FITC | Biolegend | 104505 |
| Ly6G | 1A8 | FITC | Biolegend | 127605 |
| CD45R/B220 | RA3-6B2 | FITC | Biolegend | 103206 |
| phospho-AMPK | PK136 | FITC | ThermoFischer | BS-5551R |
| PPARγ | polyclonal | FITC | ThermoFischer | BS-0530R |
| goat anti rabbit | polyclonal | AF488 | Invitrogen | A32731 |
| CD62L | MEL-14 | AF488 | Biolegend | 104420 |
| FoxP3 | 150D | AF488 | Biolegend | 320012 |
| Blimp1 | 5E7 | AF488 | Invitrogen | 53-9850-82 |
| IL13 | eBio13A | AF488 | Invitrogen | 53-7133-82 |
| CD45 | 30-F11 | BV570 | Biolegend | 103135 |
| FoxP3 | FJK-16s | PE | Invitrogen | 12-5773-80 |
| PD-1 | 29F.1A12 | PE | Biolegend | 135206 |
| phospho-mTOR | MRRBY | PE | Invitrogen | 12-9718-42 |
| I-A/I-E | M5/114.15.2 | PE | Biolegend | 107607 |
| CD103 | 2E7 | PE | Invitrogen | 12-1031-82 |
| CD36 | CRF D-2712 | PE | BD Pharmingen | 562702 |
| CD69 | H1.2F3 | PE | Biolegend | 104508 |
| IL2 | JES6-5H4 | PE | Biolegend | 503808 |
| CD45R/B220 | RA3-6B2 | PE | Biolegend | 103208 |
| Notch1 | mN1A | PE | Invitrogen | 12-5785-82 |
| Tbet | 4B10 | PE | Biolegend | 644810 |
| CD170 / SiglecF | S17007L | APC | Biolegend | 155507 |
| CD4 | RM4-5 | APC | Biolegend | 100516 |
| CD103 | 2'E7 | APC | Invitrogen | 17-1031-82 |
| TNFα | MP6-XT22 | APC | Biolegend | 506308 |
| CD8a | 53-6.7 | APC | Biolegend | 100712 |
| CD25 | PC61 | APC | Biolegend | 102012 |
| CD38 | 90 | APC | Biolegend | 102712 |
| GATA3 | 16E10A23 | APC | Biolegend | 653806 |
| CD36 | CRF D-2712 | APC | BD Pharmingen | 562744 |
| Tbet | 4B10 | PE-Cy7 | Biolegend | 644823 |
| IL4 | 11B11 | PE-Cy7 | Biolegend | 504118 |
| CD44 | IM7 | PE-Cy7 | Biolegend | 103030 |
| CD4 | GK1.5 | PE-Cy7 | Biolegend | 100422 |
| CD8a | 53-6.7 | PE-Cy7 | Invitrogen | 25-0081-82 |
| CD185/ CXCR5 | L138D7 | PE-Cy7 | Biolegend | 145516 |
| CD38 | 90 | PE-Cy7 | Biolegend | 102717 |
| TotalSeq-C0301 Hashtag 1 | M1/42; 30-F11 |  | Biolegend | 155861 |
| TotalSeq-C0301 Hashtag 2 | M1/42; 30-F11 |  | Biolegend | 155863 |
| TotalSeq-C0301 Hashtag 3 | M1/42; 30-F11 |  | Biolegend | 155865 |
| TotalSeq-C0301 Hashtag 4 | M1/42; 30-F11 |  | Biolegend | 155867 |
| TotalSeq-C0301 Hashtag 5 | M1/42; 30-F11 |  | Biolegend | 155869 |
| TotalSeq-C0301 Hashtag 6 | M1/42; 30-F11 |  | Biolegend | 155871 |
| TotalSeq™-C 0251 anti-human Hashtag 1 | LNH-94;2M |  | Biolegend | 394661 |
| TotalSeq™-C 0252 anti-human Hashtag 2 | LNH-94;2M |  | Biolegend | 394663 |
| TotalSeq™-C 0253 anti-human Hashtag 3 | LNH-94;2M |  | Biolegend | 394665 |
| TotalSeq™-C 0254 anti-human Hashtag 4 | LNH-94;2M |  | Biolegend | 394667 |
| TotalSeq™-C 0255 anti-human Hashtag 5 | LNH-94;2M |  | Biolegend | 394669 |
| CFSE (Cell trace) |  |  | Life Technologies | C34554A |
| Live/Dead |  |  | Life Technologies | L34963A |
| 2'NBDG |  | FITC | Life Technologies | N13195 |
| MitoSOX (red) |  | PE | Life Technologies | M36008 |
| MitoTracker (green) |  | FITC | Life Technologies | M7514 |
| Anti-CD69 | Polyclonal |  | ThermoFisher | PA5-114989 |
| Anti-CD4 Vio R667 | REA1211 |  | Miltenyi Biotec | 130-128-741 |
| Anti-SIRPα | P84 | Alexa Fluor 594 | BioLegend | 144020 |
| Anti-MHC II | M5/114.15.2 | Alexa Fluor 700 | ThermoFisher | 56-5321-82 |
| Anti-rabbit | Polyclonal | Alexa Fluor 546 | ThermoFisher | A32794 |
| SYTOX Green | - |  | ThermoFisher | S7020 |

| **Reagent** | **Company** | **Cat. No.** |
| --- | --- | --- |
| Paraformaldehyde | Sigma-Aldrich | P6148-1KG |
| N,N,N′,N′-Tetrakis(2-Hydroxypropyl)ethylenediamine | Sigma-Aldrich | 122262-1L |
| Dimethylsulfoxide | Sigma-Aldrich | D8418-500ML |
| Glycine, ReagentPlus®, ≥99% (HPLC) | Sigma-Aldrich | G7126-500G |
| Triton X-100 | Sigma-Aldrich | X100-500ML |
| Methanol, suitable for HPLC, ≥99.9% | Sigma-Aldrich | 34860-2.5L-R |
| Hydrogen peroxide 35% | Carl Roth GmbH + Co. KG | 9683.4 |
| Fetal Bovine Serum (FBS) | Merck KGaA, Darmstadt, Deutschland | F7524-500ML |
| Sodium azide | Sigma-Aldrich | 71289-5G |
| Heparin sodium salt | Sigma-Aldrich | H3393-25KU |
| TWEEN® 20 | Sigma-Aldrich | P1379-500ML |
| SeaKem® LE Agarose | Lonza | 50004 |
| Ethanol absolut ≥99,8% | VWR Chemicals | 20821.330P |
| Ethyl cinnamate >98% | Sigma-Aldrich | W243000-1KG-K |
