## Supplementary material for "CD36 is a metabolic checkpoint for Th2 cell tissue residency during allergic airway inflammation": Suppl. Figures

Suppl. Figure 1

A

B

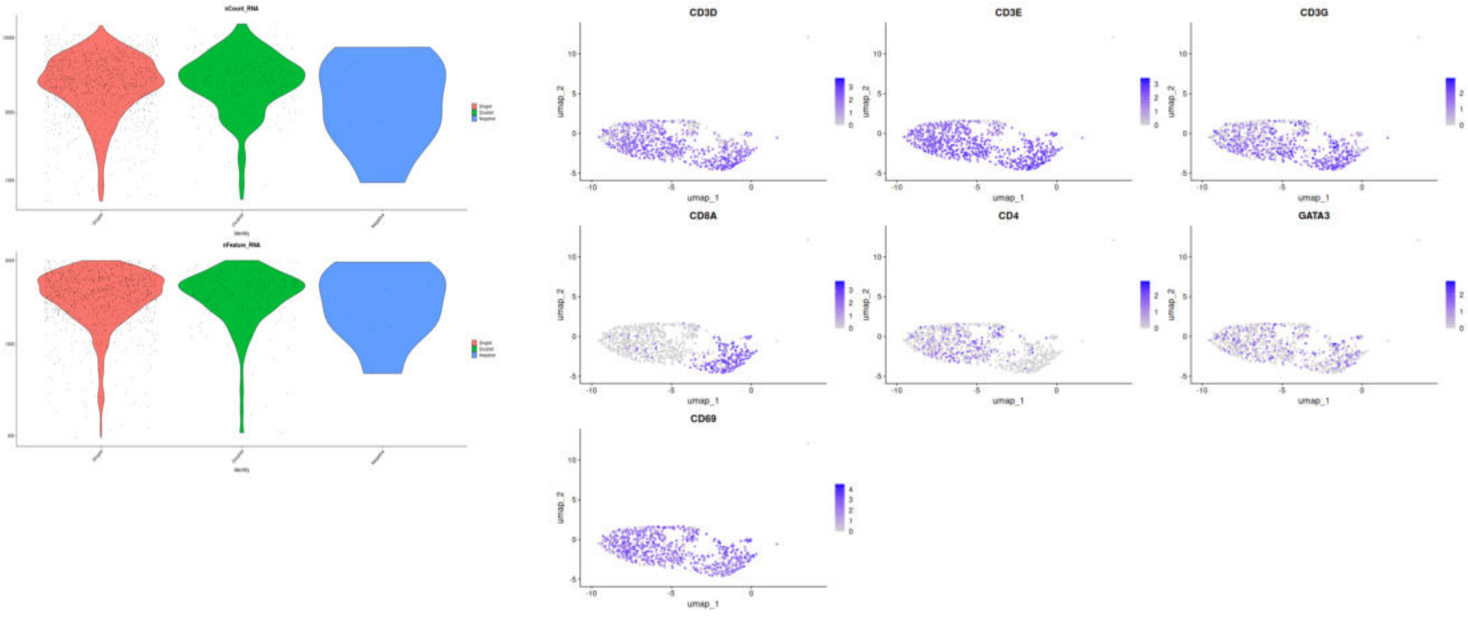

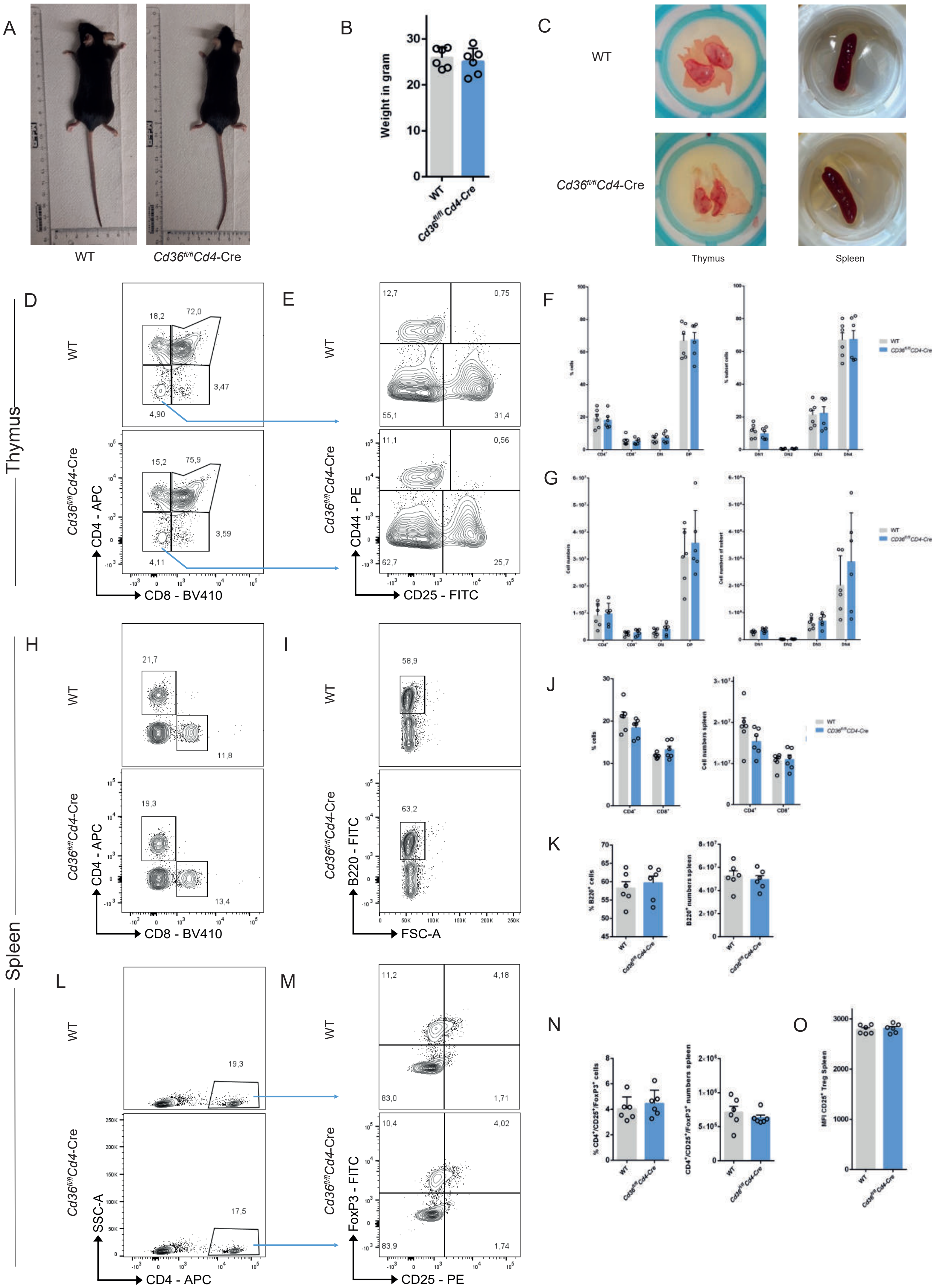
A
WT
Cd36fl/flCd4-Cre
B
Weight in gram
WT
Cd36fl/fl Cd4-Cre
C
WT
Cd36fl/flCd4-Cre
Thymus
Spleen
Thymus
D
E
F
G
CD4 - APC
CD8 - BV410
CD44 - PE
CD25 - FITC
Spleen
H
I
J
K
B220 - FITC
FSC-A
L
M
N
O
SSC-A
CD4 - APC
FoxP3 - FITC
CD25 - PE
% CD4+/CD25+/FoxP3+ cells
CD4+/CD25+/FoxP3+ numbers spleen
MFI CD25+ Treg Spleen

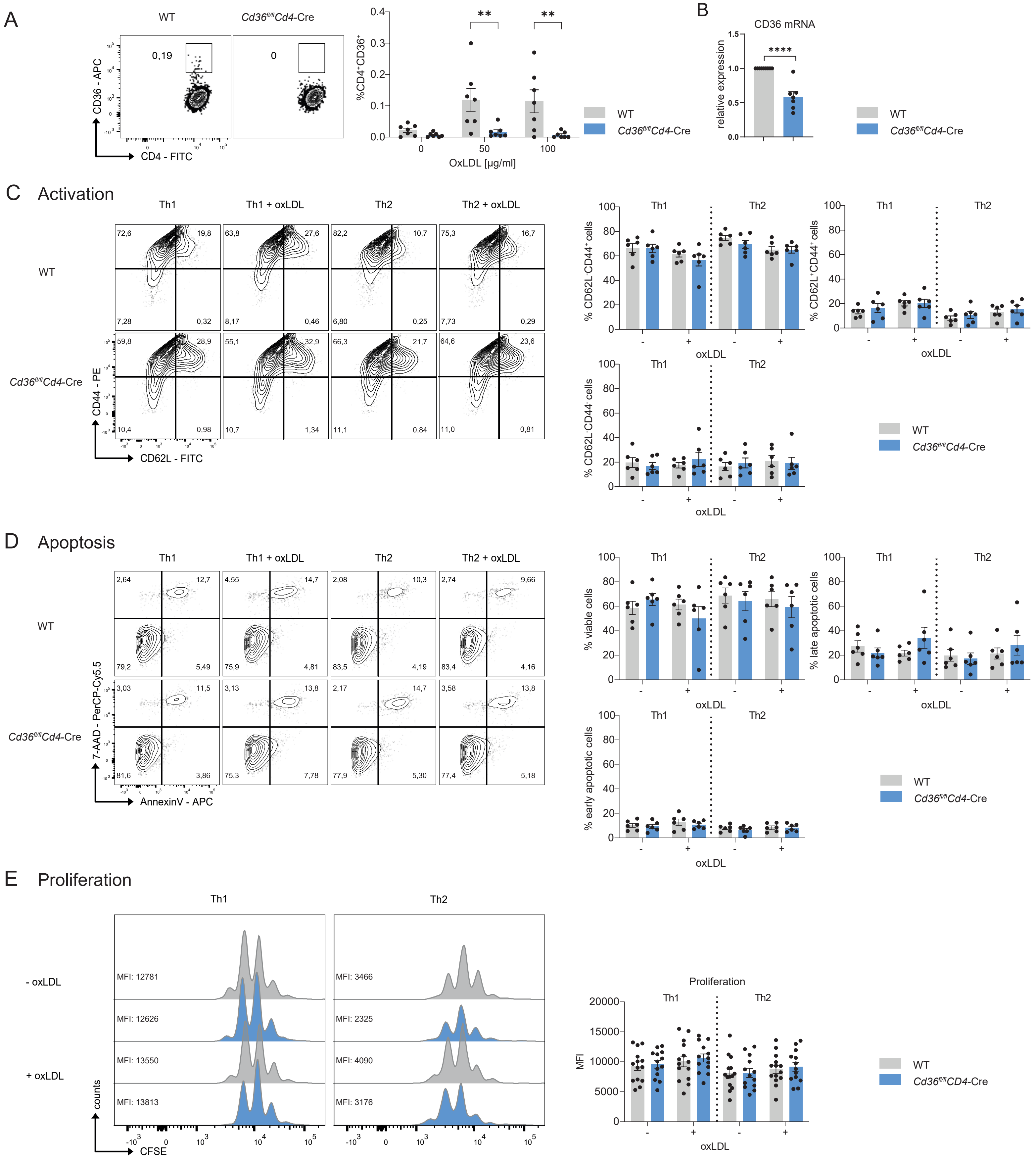
A
WT
Cd36fl/flCd4-Cre
CD36 - APC
CD4 - FITC
0,19
0
%CD4+CD36+
OxLDL [µg/ml]
B
CD36 mRNA
relative expression
C Activation
Th1
Th1 + oxLDL
Th2
Th2 + oxLDL
CD44 - PE
CD62L - FITC
% CD62L-CD44+cells
% CD62L+CD44+cells
% CD62L-CD44- cells
oxLDL
D Apoptosis
7-AAD - PerCP-Cy5.5
AnnexinV - APC
% viable cells
% late apoptotic cells
% early apoptotic cells
E Proliferation
- oxLDL
+ oxLDL
counts
CFSE
MFI: 12781
MFI: 12626
MFI: 13550
MFI: 13813
MFI: 3466
MFI: 2325
MFI: 4090
MFI: 3176
Proliferation
MFI
Cd36fl/flCD4-Cre

Suppl. Figure 4

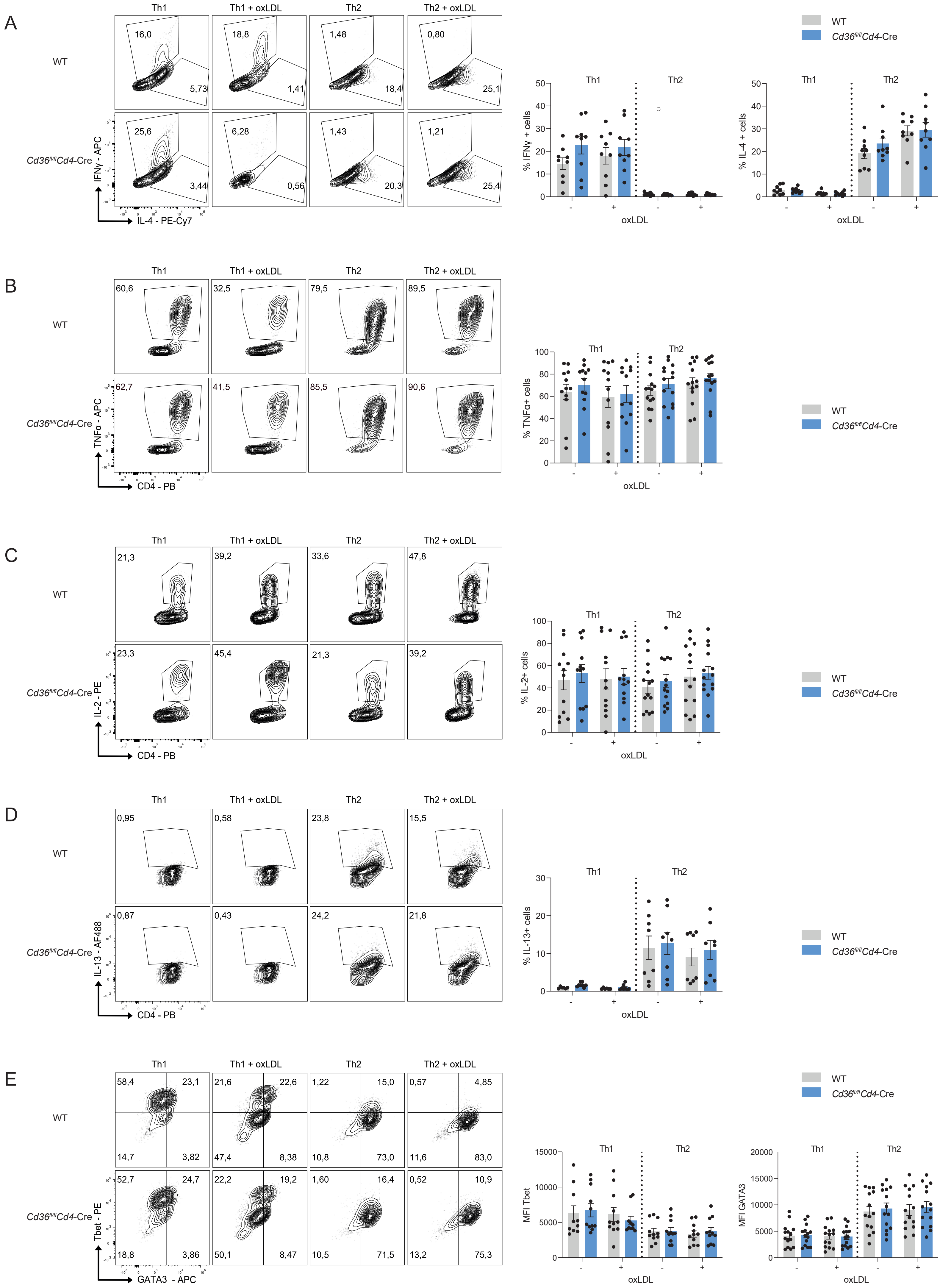

Suppl. Figure 5

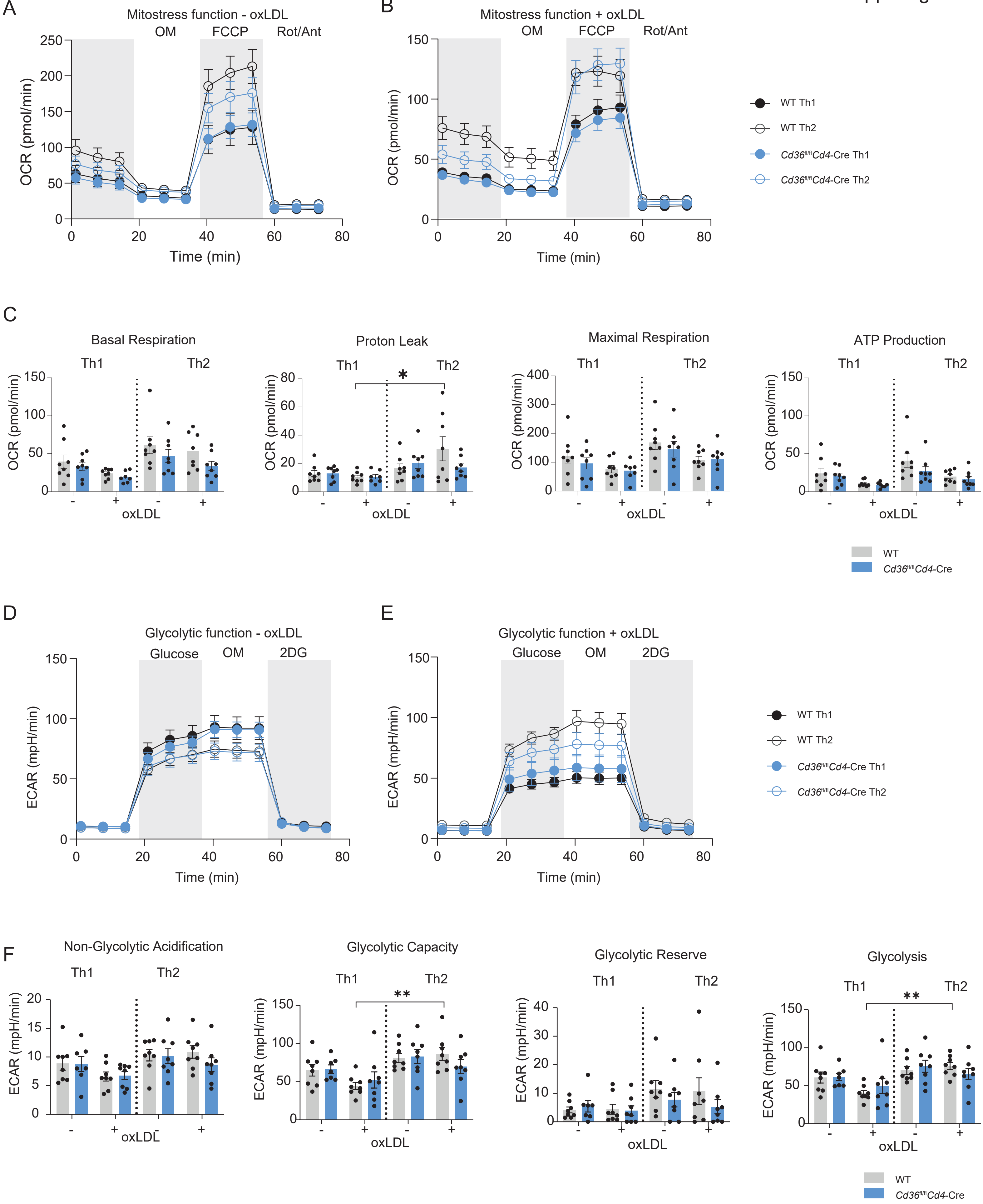

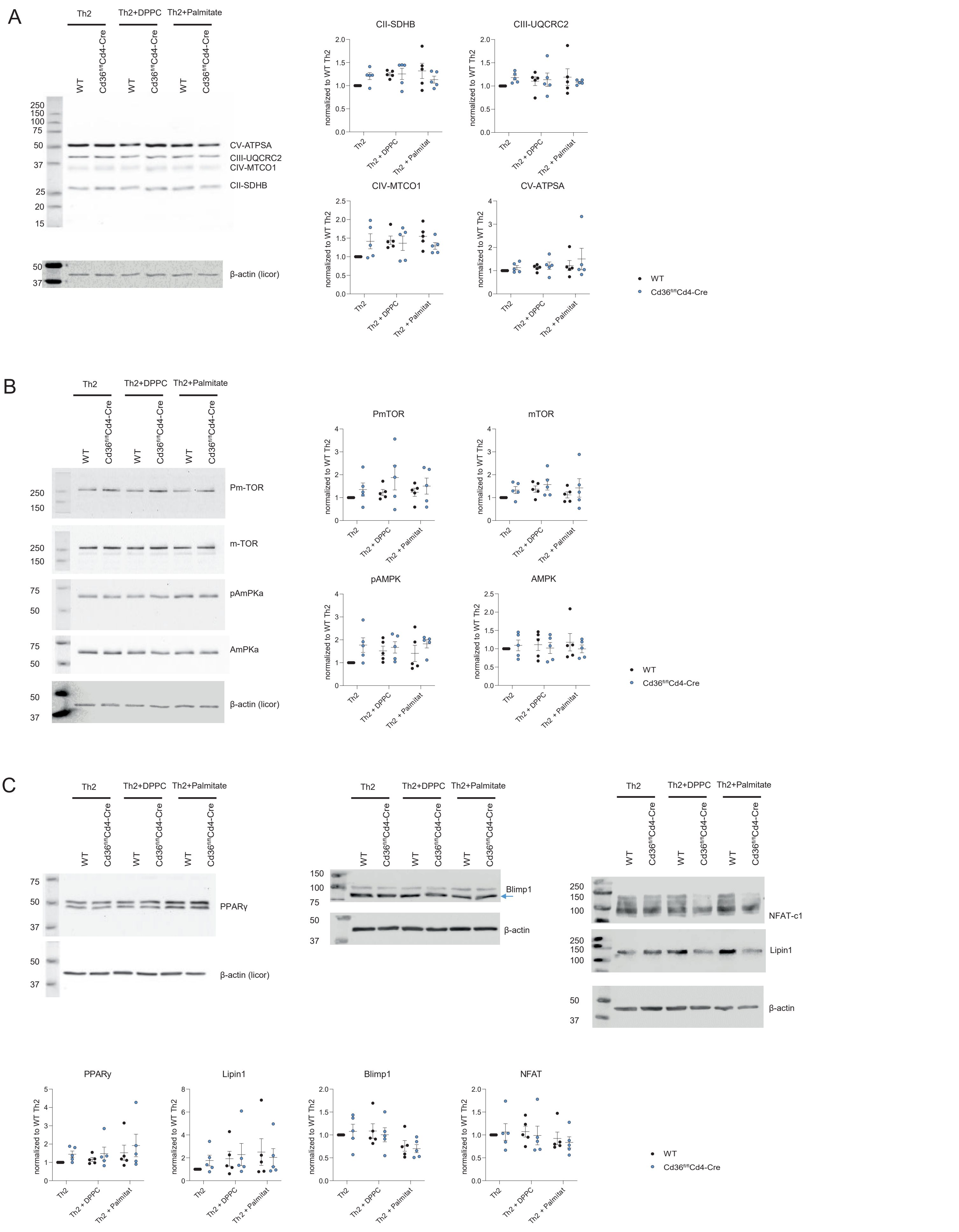
A
Th2
Th2+DPPC
Th2+Palmitate
WT
Cd36fl/flCd4-Cre
250
150
100
75
50
37
25
20
15
CV-ATPSA
CIII-UQCRC2
CIV-MTCO1
CII-SDHB
β-actin (licor)
CII-SDHB
CIII-UQCRC2
CIV-MTCO1
CV-ATPSA
normalized to WT Th2
Th2
Th2 + DPPC
Th2 + Palmitat
WT
Cd36fl/flCd4-Cre
B
Pm-TOR
m-TOR
pAmPKa
AmPKa
PmTOR
mTOR
pAMPK
AMPK
C
PPARγ
Blimp1
β-actin
NFAT-c1
Lipin1
PPARγ
Lipin1
Blimp1
NFAT

Suppl. Figure 7

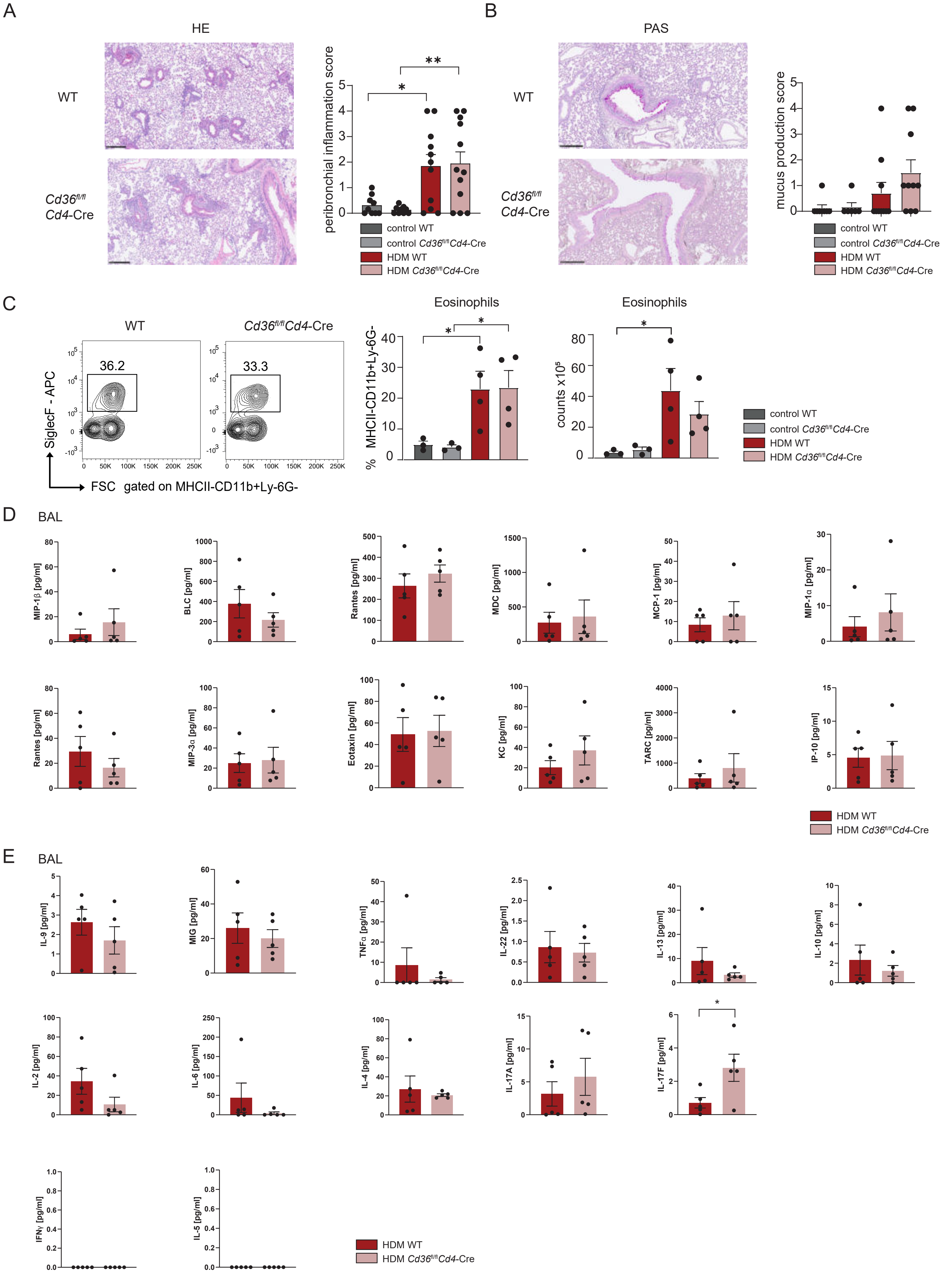

Suppl. Figure 8

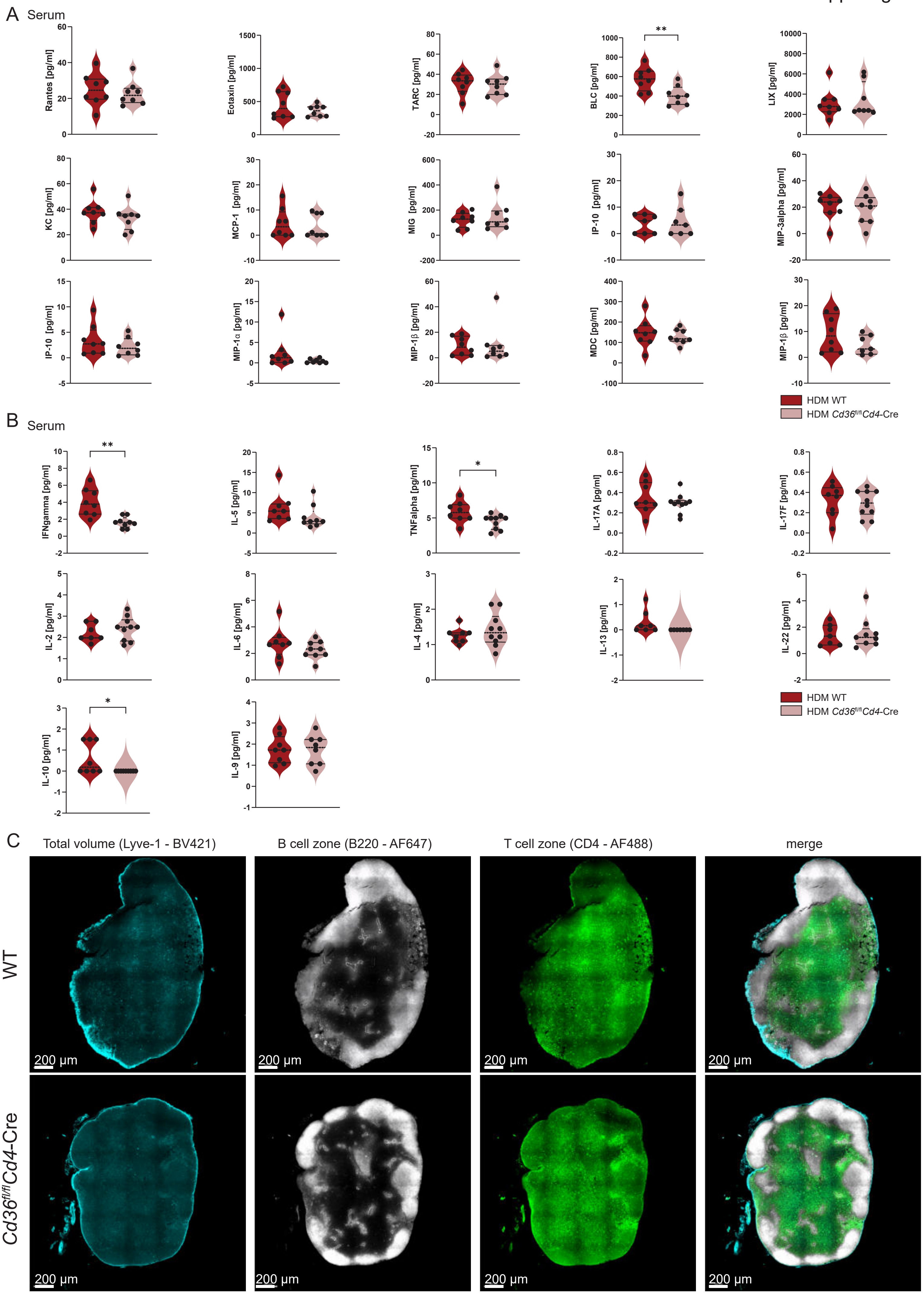

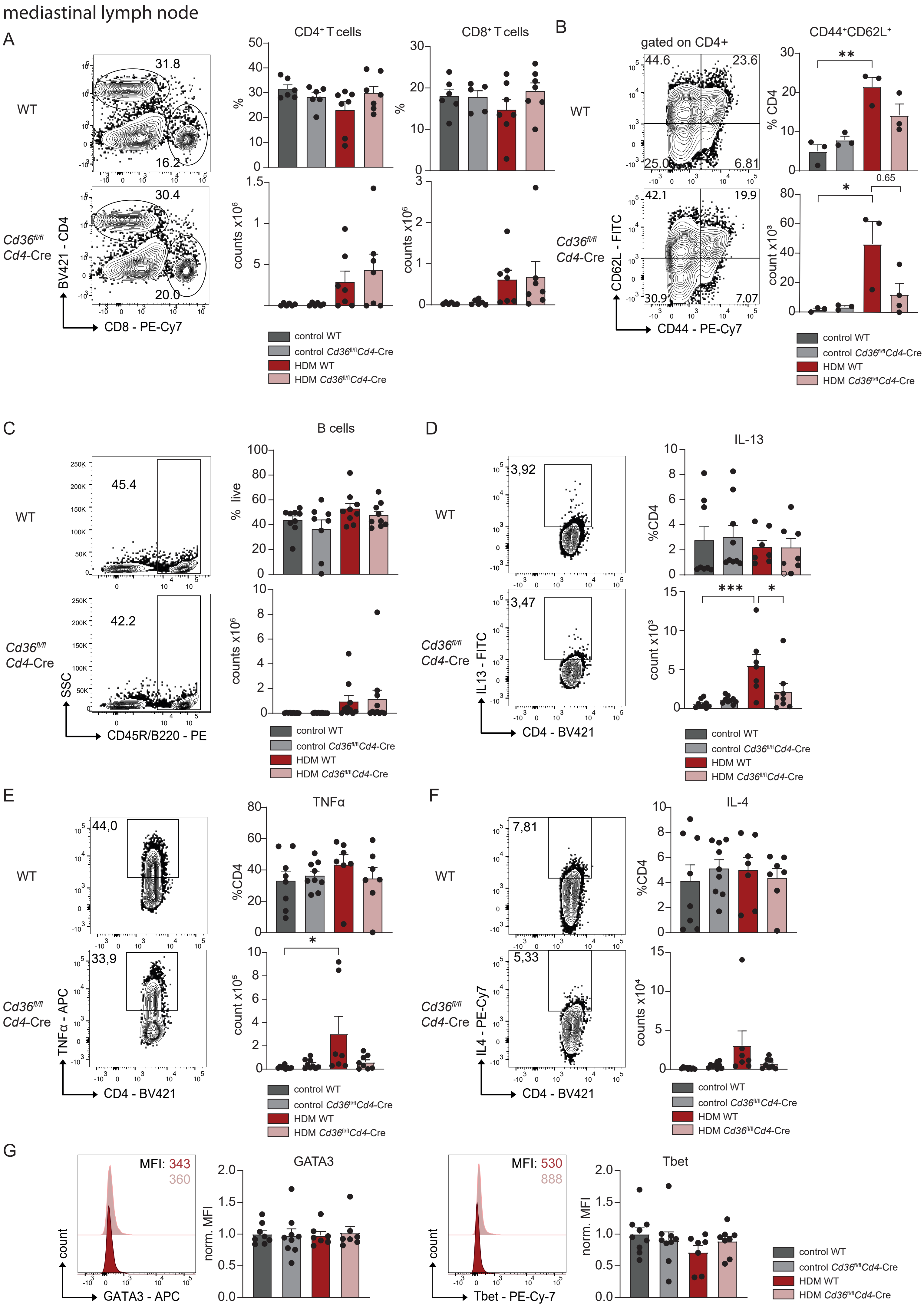
mediastinal lymph node
A
WT
Cd36fl/fl Cd4-Cre
BV421 - CD4
CD8 - PE-Cy7
31.8
16.2
30.4
20.0
CD4+ T cells
CD8+ T cells
%
counts x10^6
control WT
control Cd36fl/fl Cd4-Cre
HDM WT
HDM Cd36fl/fl Cd4-Cre
B
gated on CD4+
CD62L - FITC
CD44 - PE-Cy7
44.6
23.6
25.0
6.81
42.1
19.9
30.9
7.07
CD44+CD62L+
% CD4
count x10^3
0.65
C
B cells
SSC
CD45R/B220 - PE
45.4
42.2
% live
D
IL-13
IL13 - FITC
CD4 - BV421
3,92
3,47
%CD4
E
TNFα
TNFα - APC
44,0
33,9
count x10^5
F
IL-4
IL4 - PE-Cy7
7,81
5,33
counts x10^4
G
GATA3
Tbet
count
GATA3 - APC
Tbet - PE-Cy-7
MFI: 343
360
MFI: 530
888
norm. MFI

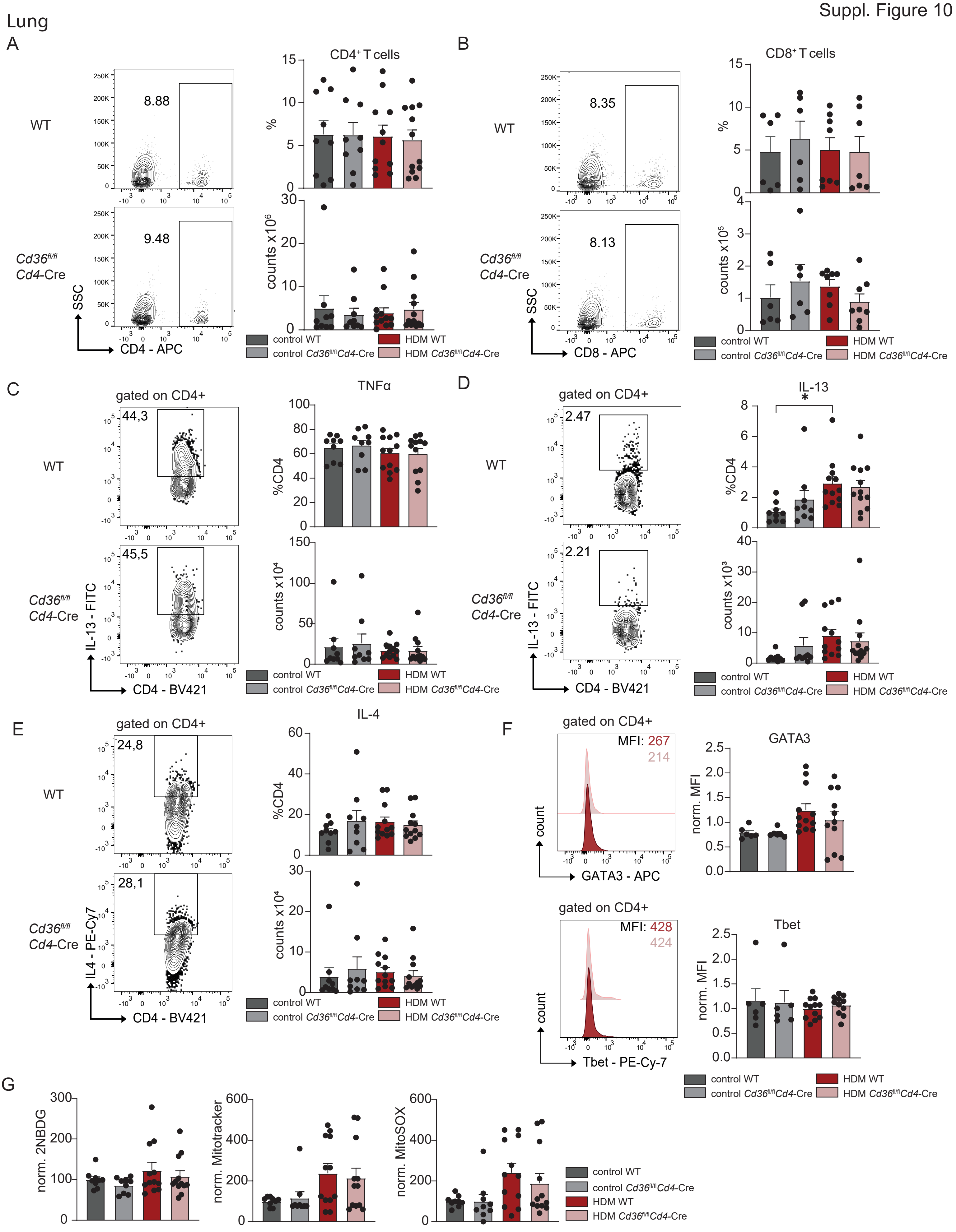
Lung
A
B
C
D
E
F
G
CD4+ T cells
CD8+ T cells
WT
Cd36fl/fl Cd4-Cre
8.88
9.48
8.35
8.13
SSC
CD4 - APC
CD8 - APC
%
counts x10^6
counts x10^5
control WT
control Cd36fl/fl Cd4-Cre
HDM WT
HDM Cd36fl/fl Cd4-Cre
gated on CD4+
TNFα
IL-13
44,3
45,5
2.47
2.21
IL-13 - FITC
CD4 - BV421
%CD4
counts x10^4
counts x10^3
IL-4
24,8
28,1
IL4 - PE-Cy7
MFI: 267
214
MFI: 428
424
count
GATA3 - APC
Tbet - PE-Cy-7
GATA3
Tbet
norm. MFI
norm. 2NBDG
norm. Mitotracker
norm. MitoSOX

Suppl. Figure 11

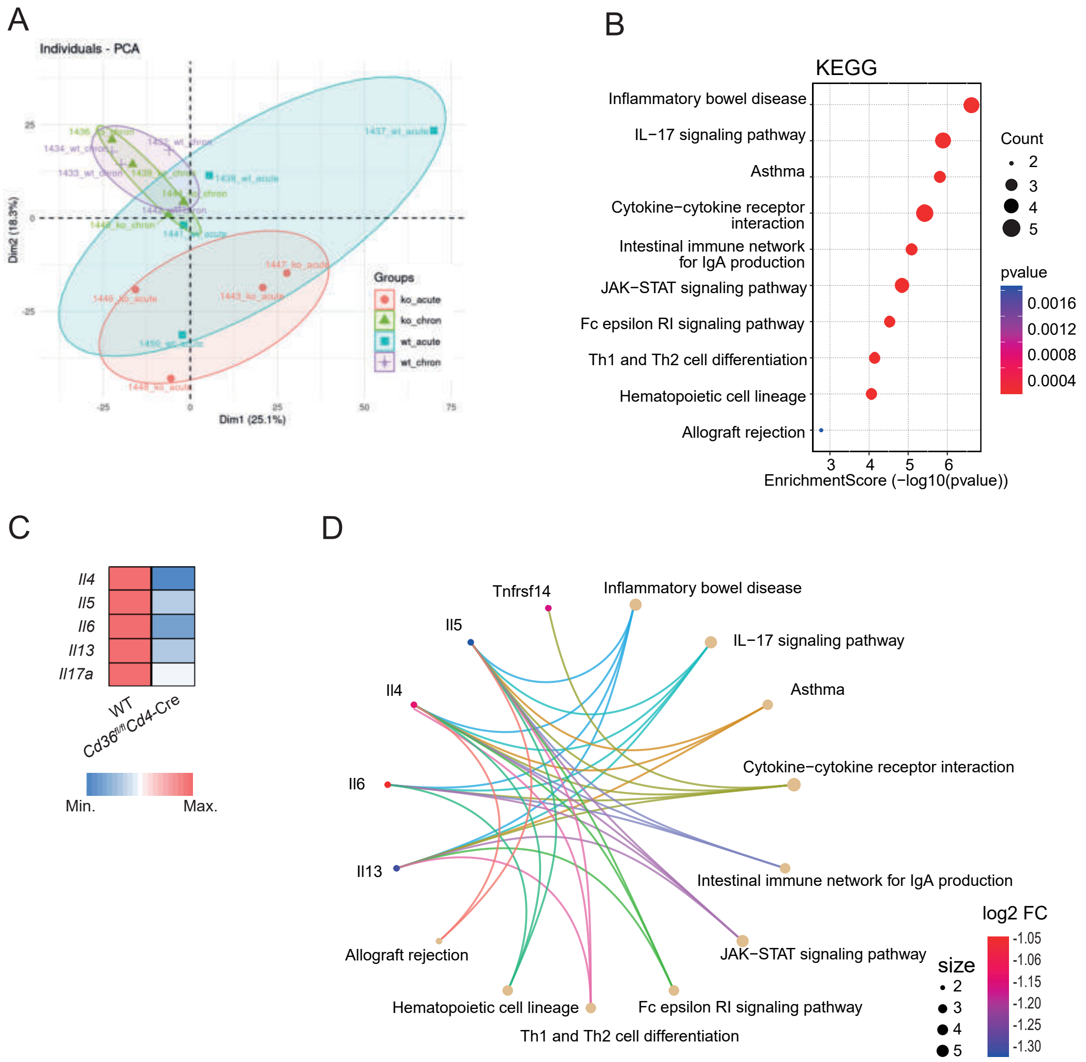

**CD36 is a metabolic checkpoint for Th2 cell tissue residency during allergic airway inflammation**

Anna Krone[1*], Simon Schreiber[1*], Nouria Jantz-Naeem[1], Anja Sammt[1], Jan Dudeck[1,2], Konstantinos Katsoulis-Dimitriou[1], Alexander Goihl[1], Oliver Hihn[6,10], Linda Black[1,3], Tobias Franz[1], Jonas Negele[1],Camilla Merten[1], Anna Marks[1], Burkhart Schraven[1,8,9],Martin Böttcher[8,11], Sarah Sandmann[13], Julian Varghese[13], Thomas Tüting[8,12], Dimitrios Mougiakakos[8,11] Jens Schreiber[3,8,9], Robert Geffers[4], Andreas Müller[1,2,8,9], Dirk Reinhold[1], Melanie Fachet[5,9], Sabine Stegemann-Koniszewski[3], Anne Dudeck[1,2,8,9], Bettina Weigelin[6,10], Sascha Kahlfuss[1,7,8,9,#]

[1] Institute of Molecular and Clinical Immunology, Medical Faculty, Otto-von-Guericke University Magdeburg, Magdeburg, Germany.

[2] Multi-Parametric Bioimaging and Cytometry Platform (MPBIC), Medical Faculty, Otto-von-Guericke University

[3] Experimental Pneumology, Department of Pneumology, University Hospital Magdeburg, Health Campus Immunology, Infectiology and Inflammation, Otto-von-Guericke University, Magdeburg, Germany.

[4] Helmholtz Center for Infection Research (HZI), Genome Analytics, Braunschweig, Germany.

[5] Institute for Medical Technology, Faculty of Electrical Engineering and Information Technology, Otto-von-Guericke University Magdeburg, Magdeburg, Germany.

[6] Department of Preclinical Imaging and Radiopharmacy, Werner Siemens Imaging Center, Eberhard Karls University, Tübingen, Germany.

[7] Institute of Medical Microbiology and Hospital Hygiene, Medical Faculty, Otto-von-Guericke University Magdeburg, Magdeburg, Germany.

[8] Health Campus Immunology, Infectiology and Inflammation (GCI[3]), Medical Faculty, Otto-von-Guericke University Magdeburg, Magdeburg, Germany.

[9] Center for Health and Medical Prevention (CHaMP), Otto-von-Guericke-University Magdeburg, Magdeburg, Germany.

[10] Cluster of Excellence iFIT (EXC 2180) "Image-Guided and Functionally Instructed Tumor Therapies", Eberhard Karls University, Tübingen, Germany.

[11] Department of Hematology and Oncology, University Hospital Magdeburg, Otto-von-Guericke University Magdeburg, Magdeburg, Germany.

[12] Experimental Dermatology, Department of Dermatology, Otto-von-Guericke University, Magdeburg, Germany.

[13] Institute of Medical Data Science, Otto-von-Guericke University, Magdeburg, Germany.

* Contributed equally

**Suppl. Figure 1: Unbiased analysis of sputum CD4$^+$ T cells using scRNA-Seq and assessment of volatile organic compounds (VOCs) reveal significant differences in lipid metabolism in asthma.**

(A) Violin plots showing gene count and feature for all singlets, doublet and hashtag negative cells for all detected cells after quality cut off.
(B) Feature plots showing hallmark genes in T cell cluster.

Sample sizes: (A)-(B) HD (n = 3). Statistical significance determined using GLM multifactorial ANOVA: *p < 0.05, **p < 0.01, ***p < 0.001.

**Suppl. Figure 2: *Cd36$^{fl/fl}$Cd4*-Cre mice show a similar appearance, body weight, and cellular composition in thymus and spleen compared to their WT littermates**

(A) Representative images of Cd36fl/flCd4-Cre and WT mice, illustrating the body size.
(B) Bodyweight 8-12 week old mice.
(C) Representative pictures of thymus and spleen.
(D) and (E) Representative dot plots illustrating the expression of CD25 and CD44 on CD4$^+$ and CD8$^+$ T cells during thymic development, highlighting the distinct double negative (DN) stages: DN1, DN2, DN3, and DN4.
(F) Graphs showing the frequencies (G) and cell numbers of CD4$^+$, CD8$^+$, DN1-4 and CD4$^+$CD8$^+$ double positive (DP) cells in the thymus.
(H) Representative dot plots of CD4$^+$ and CD8$^+$ T cells in the spleen.
(I) Representative dot plots of B220+ B cells in the spleen.
(J) Graphs showing frequencies and cell numbers of CD4$^+$ and CD8$^+$ cells in the spleen.
(K) Graphs showing frequencies and cell numbers of B cells in the spleen.
(L) And (M) Representative dot plots of regulatory T cells in the spleen.
(N) Graphs showing frequencies and cell numbers of regulatory T cells in the spleen.
(O) Graph showing CD25 expression in regulatory T cells in the spleen.

Sample sizes: n = 6; Statistical analysis was performed using two-way ANOVA followed by Šidák test for multiple comparisons (F, G, J) or Students T-Test (K, N, O); *p<0,05, **p < 0.01, ***p < 0.001.

**Suppl. Figure 3: CD36 deletion does not affect the function of Th1 and Th2 cells stimulated for 48 hours in presence or absence of 50 µg/ml oxLDL *in vitro***

(A) The bar graph shows expression of CD36 in CD4+ T cells in presence of 0, 50 and 100 µg/ml oxLDL analyzed by flow cytometry.
(B) Relative mRNA expression of *CD36*, as determined by RT-qPCR. Values are normalized to *HRPT1*.
(C) Representative dot plots and graphs showing CD62L and CD44 expression in CD4+ T cells.
(D) The apoptosis in CD4$^+$ T cells were analyzed by AnnexinV and 7AAD staining using flow cytometry. Representative dot plots and graphs display the frequencies of live cells

(AnnexinV⁻/7AAD⁻), early apoptotic cells (AnnexinV⁺/7AAD⁻), and late apoptotic cells (AnnexinV⁺/7AAD⁺).

(E) $CD4^+$ T cell proliferation was assessed by CFSE dilution assay and analyzed via flow cytometry. Representative histograms and graphs illustrating CFSE dilution in $CD4^+$ T cells.

Sample sizes: (A)-(B) n = 7; (C)-(D) n = 6; (E) n = 13-14. Statistical analysis was performed using two-way ANOVA followed by Šidák test for multiple comparisons; *p<0,05, **p < 0.01, ***p < 0.001.

**Suppl. Figure 4: CD36 deletion does not affect cytokine production or expression of transcription factor of Th1 and Th2 cells stimulated for 72 hours in presence or absence of 50 µg/ml oxLDL *in vitro***

(A) Representative dot plots and graphs showing the frequencies of IFNγ and IL-4 producing T cells.
(B) Representative dot plots and graph showing the frequencies of TNFα producing T cells.
(C) Representative dot plots and graph showing the frequencies of IL-2 producing T cells.
(D) Representative dot plots and graph showing the frequencies of IL-13 producing T cells.
(E) Representative dot plots and graphs showing the expression of GATA3 and Tbet in T cells.

Sample sizes: (A) n = 9; (B) n = 14; (C) n = 12-14; (D) n = 8; (E) n = 11-14. Statistical analysis was performed using two-way ANOVA followed by Šidák test for multiple comparisons; *p<0,05, **p < 0.01, ***p < 0.001.

**Suppl. Figure 5: CD36 deletion does not impair oxidative phosphorylation and glycolytic activity of Th1 and Th2 cells activated for 72 hours in the presence or absence of 50 µg/ml oxLDL in vitro.**

(A) Oxygen consumption rate (OCR) was measured following sequential injection of oligomycin, FCCP, and rotenone/antimycin A in absence or (B) presence of oxLDL. (C) Basal respiration, proton leak, maximal respiration, and ATP production was calculated from OCR and is presented in a bar graph.
(D) Extracellular acidification rate (ECAR) was determined after sequential addition of glucose, oligomycin, and 2-deoxy-D-glucose (2-DG) in absence or (E) presence of oxLDL. (F) Non-glycolytic acidification, glycolytic capacity, glycolytic reserve, and glycolysis, derived from ECAR measurements, is shown as a bar graph.

Sample sizes: n = 8. Statistical analysis was performed using two-way ANOVA followed by Šidák test for multiple comparisons; *p<0,05, **p < 0.01, ***p < 0.001.

**Suppl. Figure 6: CD36 deletion does not impair the expression of proteins of the electron transport chain (ETC) or upstream metabolic regulator proteins of Th2 cells activated for 72 hours in the presence or absence of Dipalmitoylphosphatidylcholin (DPPC) or palmitate *in vitro*.**

(A) Representative western blot and graphs showing four complexes of the ETA normalized to β-actin and to WT Th2 cells.

(B) Representative western blot and graphs showing pAMPK, AMPK, pm-TOR and m-TOR normalized to β-actin and to WT Th2 cells.
(C) Representative western blot and graphs showing PPARγ, Lipin1, Blimp1 and NFAT normalized to β-actin and to WT Th2 cells.

Sample sizes: n = 5. Statistical analysis was performed using two-way ANOVA followed by Šidák test for multiple comparisons; *p<0,05, **p < 0.01, ***p < 0.001.

**Suppl. Figure 7: CD36 deletion does not impair Th2 house dust mite-induced asthmatic airway inflammation.**
Lungs and serum from mice of the HDM asthma model were analyzed on day 14.
(A) Peribronchial inflammation was assessed by hematoxylin and eosin (H&E) staining.
(B) Mucus production was evaluated by periodic acid-Schiff (PAS) staining of lung tissue.
(C) Eosinophil frequencies and counts in the lungs were measured by flow cytometry (n=4 mice).
(D) Chemokines and (E) cytokines in bronchoalveolar lavage (BAL) were measured by flow cytometry (n=8).

Sample sizes: (A) n = 9-12; (B) n = 8-10; (C) n = 3-4; (D)-(E) n = 8. Statistical analysis was performed using (A)-(C) one-way ANOVA followed by Tukey's test for multiple comparisons and (D)-(E) Student's T-test; *p<0,05, **p < 0.01, ***p < 0.001.

**Suppl. Figure 8: CD36 deletion does not impair chemokine or cytokine levels in BAL of mice upon house dust mite-induced asthmatic airway inflammation.**
BAL fluid and mediastinal lymph nodes from mice in the HDM-induced asthma model were analyzed on day 14.
(A) Chemokines and (B) cytokines in the serum were measured by flow cytometry.
(C) Midsection analysis of LN from *Cd36*$^{fl/f}$ *Cd4*-Cre and control mice was performed in Imaris software. B220 was used for staining the B cell zones, CD4 for T cell zones and B220, CD4 and Lyve-1 for the total LN volume.

Sample sizes: (A)-(B) n = 5. Statistical analysis was performed using Student's T-test; *p<0,05, **p < 0.01, ***p < 0.001.

**Suppl. Figure 9: Effect of the CD36 deletion in mediastinal lymph nodes in house dust mite-induced asthmatic airway inflammation.**
Mediastinal lymph nodes were collected on day 14 from mice subjected to the HDM-induced asthma model and analyzed using flow cytometry. Representative dot plots and graphs showing frequencies and counts of
(A) $CD4^+$ and $CD8^+$ T cells,
(B) $CD4^+CD44^+CD62L^+$ T cells,
(C) B cells,
(D) IL-13 producing $CD4^+$ T cells,
(E) TNFα producing $CD4^+$ T cells, and
(F) IL-4 producing $CD4^+$ T cells.
(G) Representative histograms and mean fluorescence intensity (MFI) of GATA3 and T-bet expression in $CD4^+$ T cells.

Sample sizes: (A) n = 5-7; (B) n = 3; (C) n = 9-10; (D)-(G) n = 7-9. Statistical analysis was performed using one-way ANOVA followed by Tukey's test for multiple comparisons; *p<0,05, **p < 0.01, ***p < 0.001.

**Suppl. Figure 10: Effect of the CD36 deletion in lungs in house dust mite-induced asthmatic airway inflammation.**
Lungs were collected on day 14 from mice subjected to the HDM-induced asthma model and analyzed using flow cytometry. Representative dotplots and graphs showing frequencies and counts of

(A) $CD4^+$ T cells,
(B) $CD8^+$ T cells
(C) TNFα producing $CD4^+$ T cells,
(D) IL-13 producing $CD4^+$ T cells, and
(E) IL-4 producing $CD4^+$ T cells.
(F) Representative histograms and mean fluorescence intensity (MFI) of GATA3 and T-bet expression in $CD4^+$ T cells.
(G) Glucose uptake was evaluated by the uptake of 2-NBDG, mitochondrial content was evaluated by MitoTracker MFI, and mitochondrial ROS were quantified by MitoSOX MFI using flow cytometry.

Sample sizes: (A) n = 9-12; (B) n = 6-8; (C)-(H) n = 9-12; (D). Statistical analysis was performed using one-way ANOVA followed by Tukey's test for multiple comparisons; *p<0,05, **p < 0.01, ***p < 0.001.

**Suppl. Figure 11: Analysis of WT versus *Cd36* $^{fl/fl}$*Cd4*-Cre mice upon induction of chronic HDM allergic airway inflammation**

(A) Principal Component Plot (PCA) plot shows clusters of sample similarity of three wild-type and three *Cd36*$^{fl/fl}$*Cd4*-Cre mice upon induction of acute and chronic HDM allergic airway inflammation.
(B) KEGG pathway enrichment analysis of DEGs of wild-type versus *Cd36*$^{fl/fl}$*Cd4*-Cre mice upon induction of chronic HDM allergic airway inflammation Log2FC > 1. p-value<0.05
(C) Heatmap of normalized counts of selected, asthmatic genes of wild-type versus *Cd36*$^{fl/fl}$*Cd4*-Cre mice upon induction of chronic HDM allergic airway inflammation.
(D) Netplot showing connection of DEGs related to their enriched pathways.

Sample sizes: n = 3
